# Tumor-Immune Partitioning and Clustering (TIPC) algorithm reveals distinct signatures of tumor-immune cell interactions within the tumor microenvironment

**DOI:** 10.1101/2020.05.29.111542

**Authors:** Mai Chan Lau, Jennifer Borowsky, Juha P. Väyrynen, Koichiro Haruki, Melissa Zhao, Andressa Dias Costa, Simeng Gu, Annacarolina da Silva, Kota Arima, Joe Yeong, Kristen D. Felt, Tsuyoshi Hamada, Reiko Nishihara, Jochen K. Lennerz, Charles S. Fuchs, Catherine J. Wu, Shuji Ogino, Jonathan A. Nowak

## Abstract

Growing evidence supports the importance of understanding tumor-immune spatial relationship in the tumor microenvironment in order to achieve precision cancer therapy. However, existing methods, based on oversimplistic cell-to-cell proximity, are largely confounded by immune cell density and are ineffective in capturing tumor-immune spatial patterns. Here we developed a novel computational algorithm, termed Tumor-Immune Partitioning and Clustering (TIPC), to offer an effective solution for spatially informed tumor subtyping. Our method could measure the extent of immune cell partitioning between tumor epithelial and stromal areas as well as the degree of immune cell clustering. Using a U.S. nation-wide colorectal cancer database, we showed that TIPC could determine tumor subtypes with unique tumor-immune spatial patterns that were significantly associated with patient survival and key tumor molecular features. We also demonstrated that TIPC was robust to parameter settings and readily applicable to different immune cell types. The capability of TIPC in delineating clinically relevant patient subtypes that encapsulate tumor-immune spatial relationship, immune density, and tumor morphology is expected to shed light on underlying immune mechanisms. Hence, TIPC can be a useful bioinformatics tool for effective characterization of the spatial composition of the tumor-immune microenvironment to inform precision immunotherapy.

## Introduction

Increasing evidence suggests that not only immune cell abundance but also the spatial relationship between immune and tumor cells play a crucial role in cancer development, progression, treatment responsiveness, and clinical outcome^1–3^. Comprehensive spatial and phenotypic characterization of immune cells in solid tumors is vastly empowered by the recent developments in tissue-based multiplexing techniques including multiplex immunofluorescence (mIF)^4–7^, imaging mass cytometry (IMC)^8, 9^, multiplexed ion beam imaging (MIBI)^10, 11^. Along with the augmented ability of advanced digital image processing techniques, it enables accurate tissue and cell segmentation leading to better appreciation of tumor-immune interaction at single-cell resolution.

While the literature has been largely limited to immune cell density-based analysis, there is a growing interest in understanding the tumor-immune spatial heterogeneity in tumor microenvironment (TME)^12–14^. Existing approaches include direct cell distance measurements as well as more sophisticated computational methods: (1) Morisita-Horn (MH) index, an ecological measure adopted to quantify the co-localization between immune and tumor cells^15^; (2) methods derived from spatial point patterns theory, including G-cross^16^ and L-cross functions^12^, which estimate relative proximity between two cell types.

Although the studies that applied these approaches have found tumor subtypes with significant association with patient survival or disease recurrence, these methods have two major drawbacks. First, the tumor subtypes are largely confounded by immune densities which are often associated with patient outcomes. Second, highly heterogeneous tumor-immune spatial relationship (Supplementary Table 1) within a TME is compressed into a single numerical value, resulting in information loss. The latter is particularly detrimental to the study of solid tumors whose immune cell spatial organization is known to be confounded by tumor morphology. Fig. 1 demonstrates two simulated scenarios where MH indices and G-cross functions are unable to distinguish distinct tumor-immune spatial patterns.

**Figure 1.**
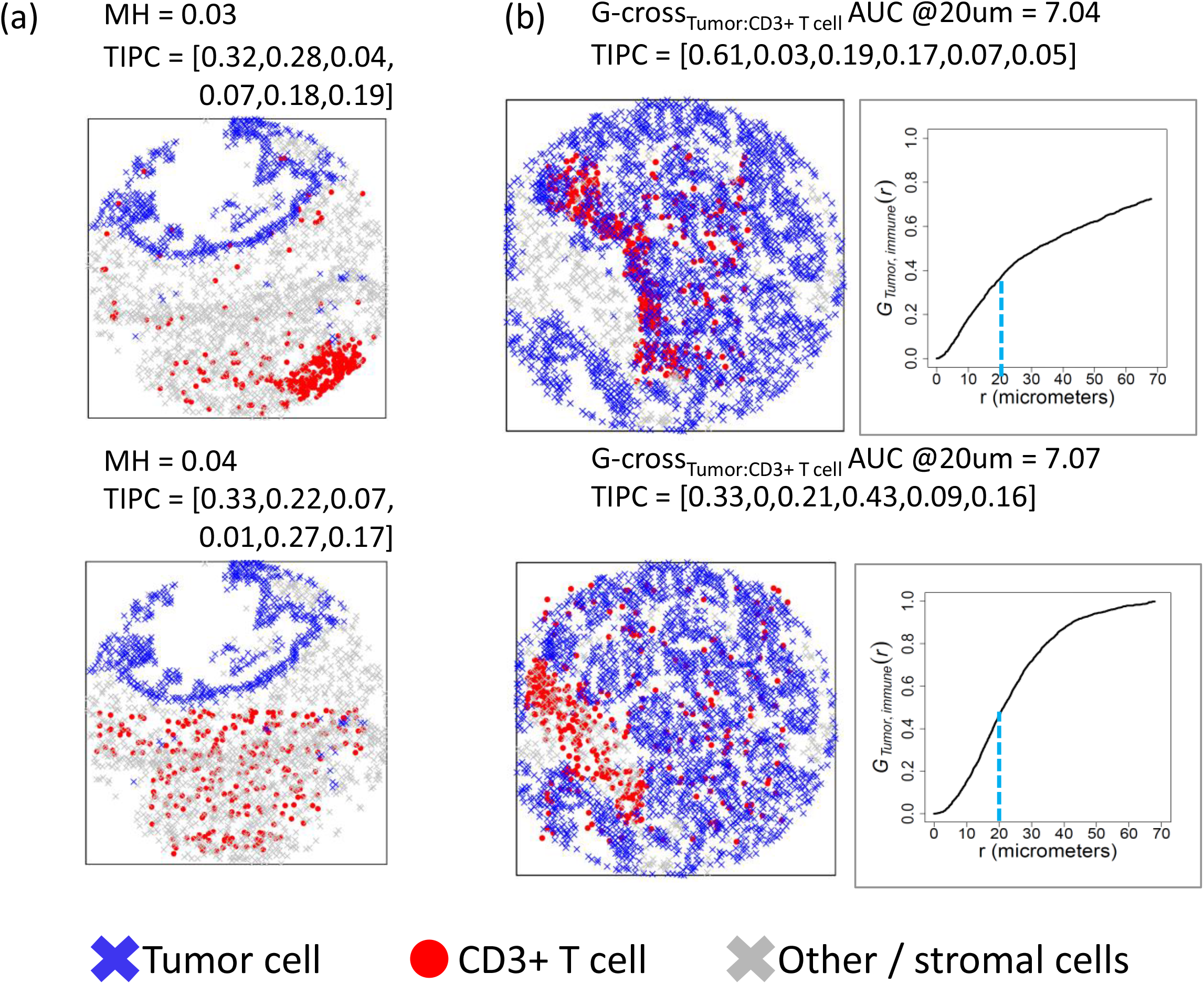
Simulated data demonstrating different spatial patterns of CD3^+^ T cell distribution in the CRC microenvironment which were, however, indistinguishable by existing methods including (**a**) Morisita-Horn (MH) index using 10-by-10 pixels rectangular grid and (**b**) G-cross function. Simulated data were generated using the same number of CD3^+^ T cells in individual examples (**a**) and (**b**). TIPC spatial parameter values, in order: tumor_ONLY, Stroma_ONLY, I2Tu_high, I2Tu_low, I2S_high, I2S_low, are computed using 70 pixels hexagonal grid.

To address these drawbacks, we propose a novel computational algorithm, termed as Tumor-Immune Partitioning and Clustering (TIPC), to offer an effective solution for spatially informed tumor subtyping in a density-agnostic manner. In this study, we first applied TIPC to a mIF T-cell data of 931 colorectal cancer (CRC) cases from two U.S. nationwide prospective cohorts, and then we validated our algorithm using neutrophils and eosinophils identified from hematoxylin and eosin (H&E) stained sections of the same tumors^17^. We also compared the performance of TIPC to existing methods namely G-cross, L-cross, and Morisita-Horn index.

## Results

TIPC is a novel computational method designed for characterizing the spatial composition of tumor-immune microenvironment. Essentially, using a hexagonal tessellation approach, TIPC simultaneously measures the degree of partitioning of immune cells in both tumor epithelial and stromal areas as well as the degree of immune cell clustering, within a TME. These measures are encapsulated concisely using six spatial parameters (Supplementary Table 2) which can be easily compared across tumors (Fig. 2). Employing an unsupervised clustering algorithm, TIPC categorizes tumors into subtypes with unique spatial patterns.

**Figure 2.**
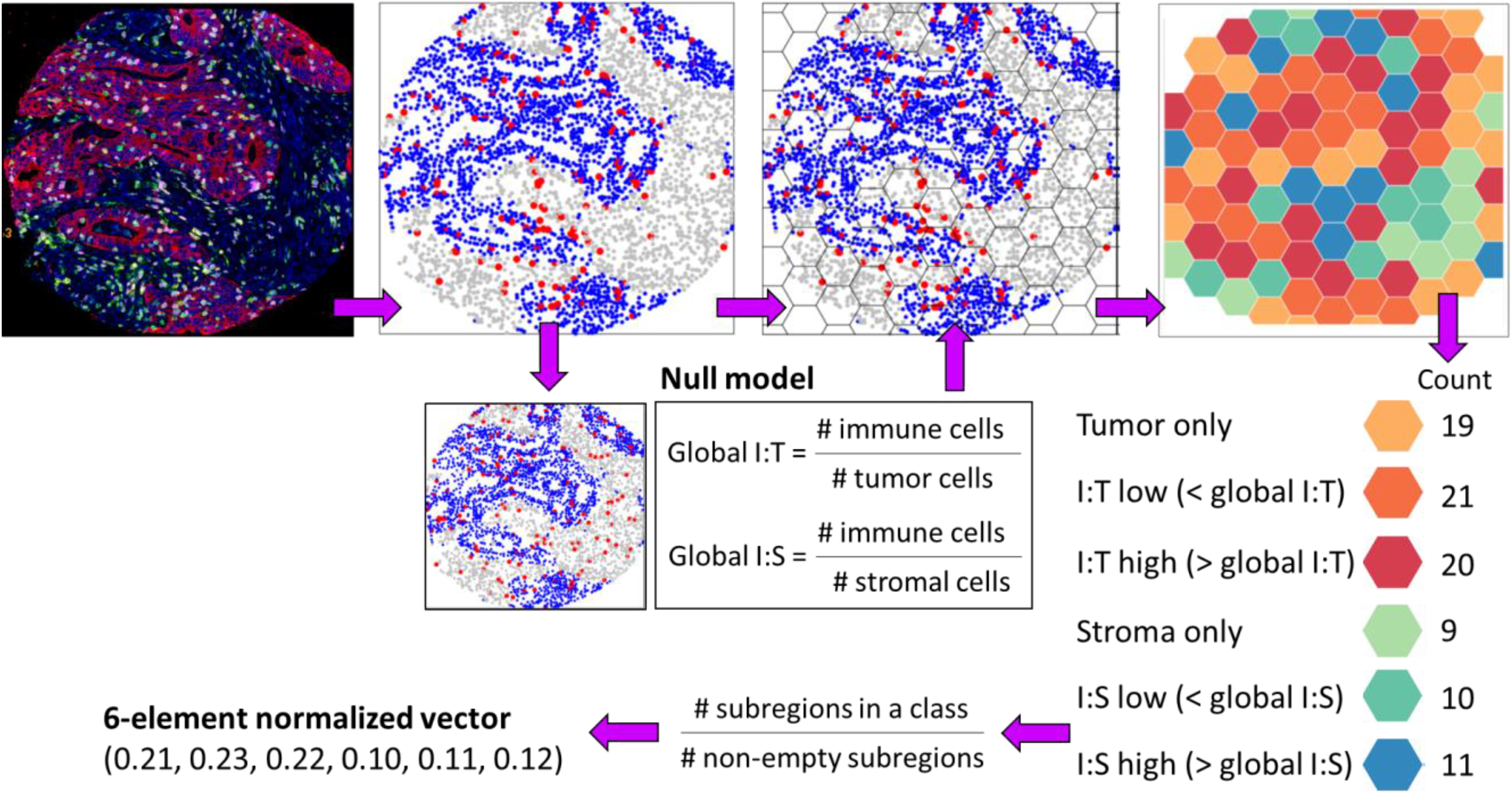
Tumor-Immune Partitioning and Clustering (TIPC) - analytic approach and implementation, using an example of multiplexed immunofluorescence stained image.

### Application of TIPC to characterize CD3^+^T-cell spatial distribution in tumor microenvironment of CRC

Here, we demonstrate determining the optimal values for two parameters involved in TIPC implementation: (1) sub-region size (*hex_len*) defining the scale of local immune spatial distribution, and (2) the number of clusters (*k*) controlling tumor subtyping resolution.

Supplementary Fig. 1 shows that as *hex_len* increased, the proportions of immune-containing sub-regions i.e., I:T high and low, I:S high and low increased, whereas the proportions of immune-absent sub-regions i.e. tumor-only and stroma-only decreased. This is because larger sub-regions have a higher tendency to contain more than one cell type. The rule of thumb is to choose a sufficiently large sub-region size to capture the information of immune partitioning between tumor epithelial and stromal areas, without over-compromising the intrinsic absence of immune cells. While both 70 and 80 pixels ensured minimal outlier tumors, the former produced more effective clusters (i.e. tumor subtypes) (Supplementary Fig. 2) and thus more favorable.

On the other hand, the deflection point in consensus cumulative distribution function (CDF) delta curve indicated there were 4 detectable tumor subtypes (Fig. 3a). The tracking plot (Fig. 3b) further reveals that stable and more granular clustering can be achieved with 9 or more clusters. To ensure sufficient statistical power, we used 9 clusters which ultimately yielded 6 major clusters with reasonable sample sizes. The clustering quality check based on heat-map visualization (Fig. 3c) was satisfactory as the 6 major clusters manifested a coherent spatial characteristics individually.

**Figure 3.**
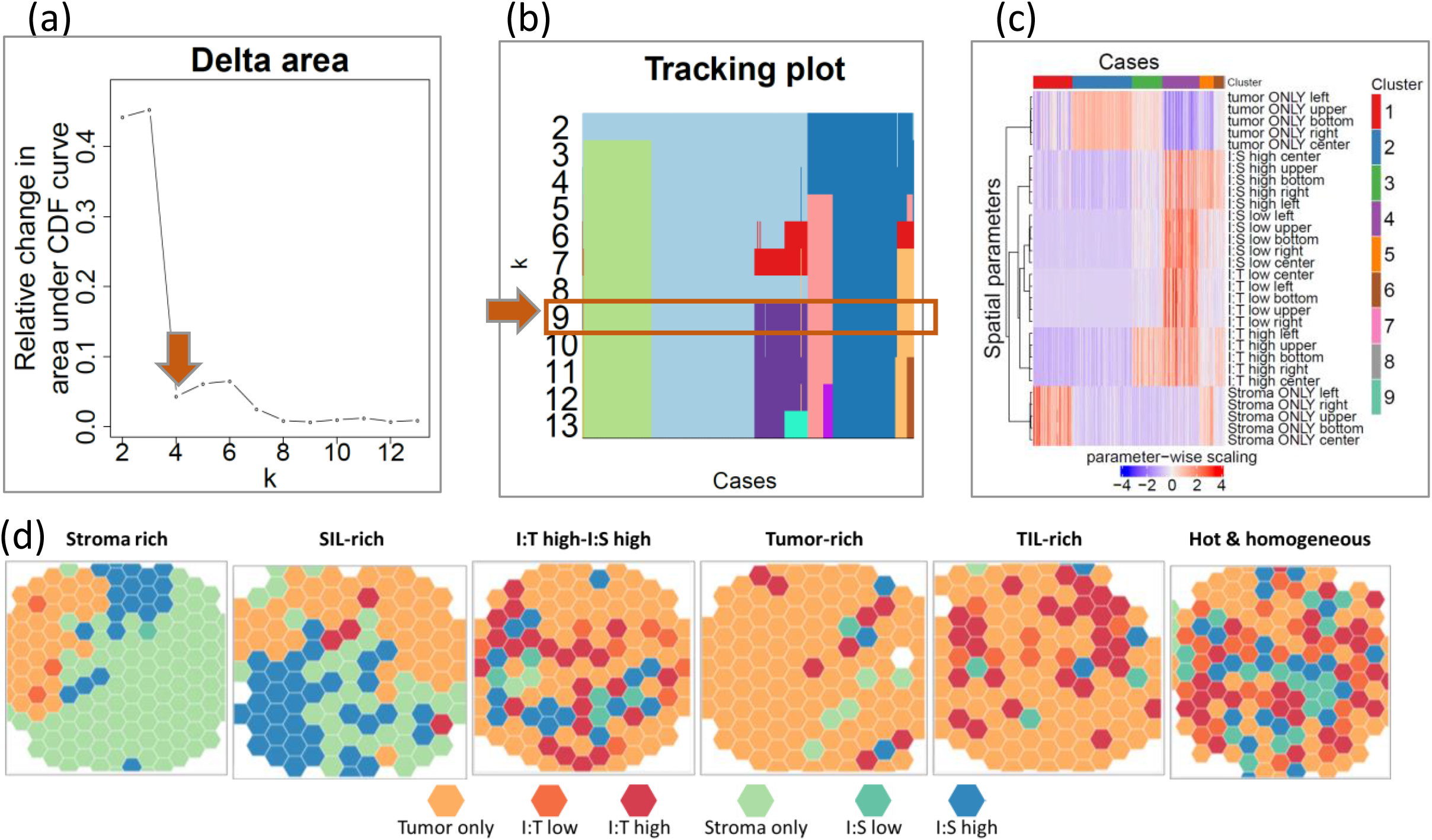
TIPC application to CRC CD3^+^ T-cell data. Selection of optimal number of clusters based on the (**a**) consensus cumulative distribution function (CDF) delta plot and (**b**) tracking plot. The resultant tumor subtypes and their spatial patterns are represented by the (**c**) heat-map. (**d**) The 6 example cases with similar CD3^+^ T-cell density (within the 3^rd^ quartile) selected from each of the 6 major tumor subtypes i.e. TIPC clusters illustrate the distinct spatial pattern, where sub-regions are color coded according to their assignment of spatial categories.

### Spatial subtypes of CD3^+^ T-cell demonstrated significant association with clinicopathologic and molecular features

Using previously selected optimal parameter values (*hex_len* = 70 pixels, *k* = 9 clusters), 927 tumors were grouped into 6 spatial subtypes with sample sizes ranging from 49 to 290, after excluding 4 tumors found in 3 outlier clusters. The 6 spatial subtypes are referred by descriptive names (Supplementary Table 3) as follows: stroma-rich, tumor-rich, TIL-rich, hot-and-homogeneous, SIL-rich, and I:T high-I:S high (Fig. 3c, read from left to right). The corresponding cluster-median T-cell densities were 19.3, 18.8, 130.2, 614.6, 199.7, and 170.1 cell/mm^2^ (Fig. 4a). In Fig. 3d, one example case was selected from each spatial subtype, with similar T-cell densities within the 3rd quartile density range, demonstrating distinct tumor-immune spatial patterns among these cases.

**Figure 4.**
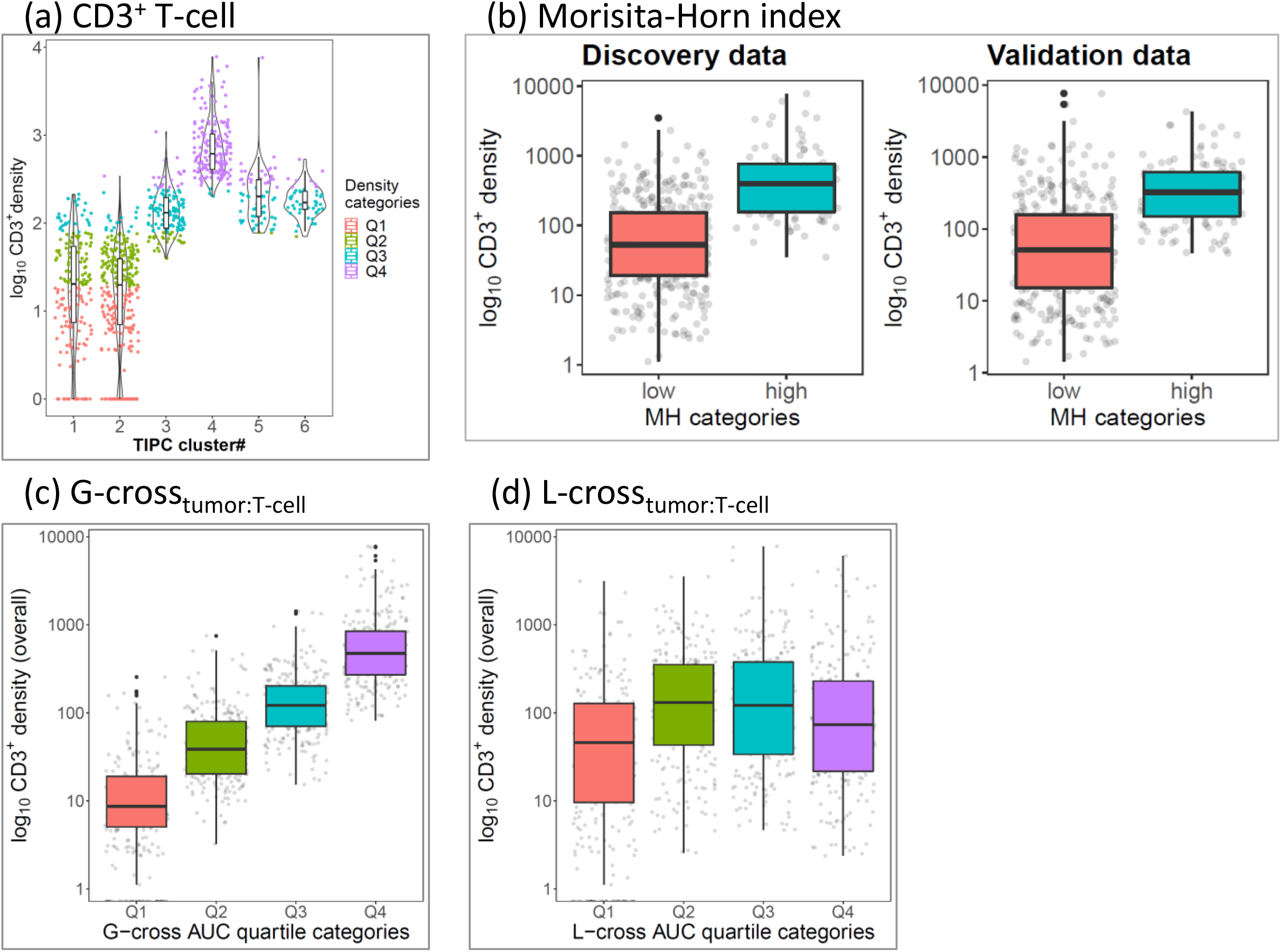
Overall CD3^+^ T-cell density distribution of tumor subtypes identified by (**a**) TIPC (*hex_len* = 70 pixels, *k* = 9) where cases were color-coded by density quartile categories, (**b**) Morisita-Horn (MH_T cell vs tumor cell_) index (left panel: discovery subset, right panel: validation subset) using 10-by-10 pixels rectangular grid and 80^th^ percentile as the cut-off; (**c**) G-cross_tumor cell vs T cell_ AUC, and (**d**) L-cross_tumor cell vs T cell_ AUC. Box plot is defined by the 25th percentile (lower) and 75th percentile (upper) while extending lines mark the minimum (lower) and maximum densities (upper), black dots represent outliers higher (or lower) than the highest (or lowest) value within 1.5× the interquartile range (IQR); jittered gray dots represent cases.

These spatial subtypes were significantly associated with 10-year CRC-specific patient outcome (log-rank p < 0.001; Fig. 5a). In both univariable and multivariable Cox proportional hazard regression analyses, using tumor-rich as the reference, the hot-and-homogeneous, I:T high-I:S high, and TIL-rich demonstrated more favorable CRC-specific survival (univariable: p < 0.001, p = 0.02, and p = 0.02; multivariable: p = 0.001, p = 0.02, and p = 0.04, respectively (Supplementary Fig. 3 shows the HRs and CIs). Interestingly, while hot-and-homogeneous (181 cases) and I:T high-I:S high (49 cases) exhibited distinct T-cell densities (the former was 3.6 times higher in median value than the latter; Fig. 4a), they demonstrated similarly favorable clinical outcome as evinced by the inseparable Kaplan-Meier curves (Fig. 5a). We verified this assertion by conducting a chi-square test which showed no significant differences between these two subtypes in terms of 5-year CRC-specific survival (p > 0.05).

**Figure 5.**
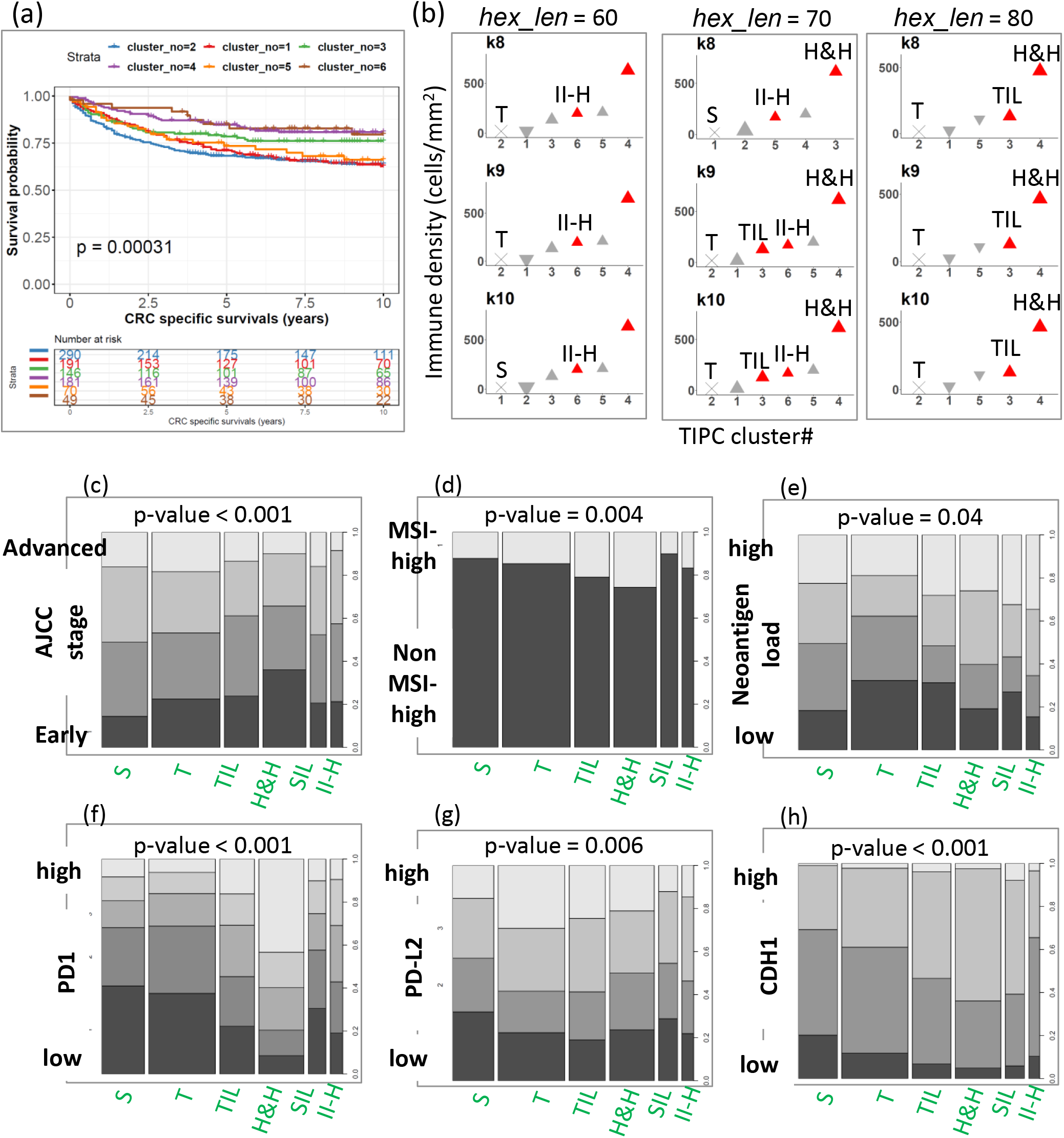
Prognostic and molecular relevance of spatial subtypes determined by TIPC using CRC CD3^+^ T-cell data. (**a**) Kaplan-Meier curves and log-rank test result. (**b**) Univariable Cox PH regression analysis result corresponding to 3 hexagon sizes (*hex_len =* 60, 70, and 90 pixels) and 3 different number of clusters (*k* = 8 to 10) where p-values < 0.05 are highlighted in red (see **Supplementary Figure 5** for full data). (**c**-**h**) 100% stacked column charts elucidate molecular compositions of each subtype on (**c**) CIMP-high status, (**d**) MSI-high status, (**e**) neoantigen load, (**f**) PDCD1 (PD1), (**g**) PDCD1LG2 (PD-L2), and (**e**) CDH1 (E-cadherin). P-values were computed using Extended Cochran–Armitage test. TIPC analysis was performed using *hex_len* = 70 pixels and *k* = 9 for (**a**), (**c**-**f**).

The Cochran-Armitage analysis revealed significant association of the tumor subtypes with tumor histologic and molecular features including the lymphocytic reactions (p < 0.001), AJCC stage (p < 0.001), and MSI-high (p = 0.004) (Fig. 5c-h, Supplementary Fig. 4). While both the hot-and-homogenous and TIL-rich tumors had nearly the same proportion of MSI-high cases, these tumors differed not only in T-cell spatial distribution but also in T-cell density, with TIL-rich tumors having a 4.7-fold lower median T-cell density, suggesting heterogeneity in the T-cell response even within MSI-high tumors. Moreover, the tumor subtypes showed a strong association with the level of tumor infiltrating PDCD1 (PD-1)-positive cells (p < 0.001) and tumor PDCD1LG2 (PD-L2) expression (p = 0.006), suggesting immune checkpoint-related mechanisms may be involved in sculpting T-cell spatial distributions. In Fig. 5h, we can see clear differences in the tumor expression of CDH1 (E-cadherin) an epithelial cell junction protein, between tumor subtypes with nearly the same T-cell density but different spatial patterns, particularly tumor-rich versus stroma-rich, and TIL-rich versus I:T high-I:S high. CDH1-loss, associated with infiltrative growth potential, was enriched in stroma-rich tumors, implying that its role in shaping tumor-immune spatial relationship.

### Comparing TIPC with existing methods using CD3^+^ T-cell data

We compared the performance of TIPC with three existing methods, namely G-cross, L-cross, and Morisita-Horn (MH) index, which have recently been adopted to study tumor-immune spatial relationship; Fig. 1 shows the comparison using simulated TME. Using CD3^+^ T-cell data, MH index was used to quantify the degree of co-localization between T-cells and tumor (or stromal) cells, with 4 different rectangular sub-region sizes of 9-by-9, 10-by-10, 11-by-11, and 12-by-12 pixels (Supplementary Fig. 6). Of note, 10-by-10 rectangular tessellation yields similar total number of sub-regions (i.e. 100) as that of hexagonal tessellation with *hex_len* of 70 pixels. Among these settings, i.e. 4 sub-region sizes each tested using 13 cut-offs equally spaced from 20th to 80th percentiles, only two showed significant survival association on both discovery and validation subsets, where tumors were dichotomized into low and high subtypes of MH_tumor:CD3+T cell_ at 80^th^ percentile. One used a 10-by-10 grid (discovery data FDR = 0.05, HR = 1.90 with 95% CI 1.13 – 3.21, and validation FDR = 0.01, HR = 1.90 with 95% CI 1.44 – 4.14) and the other used a 12-by-12 grid (discovery FDR = 0.04, HR = 1.91 with 95% CI 1.13 – 3.23, and validation FDR = 0.02, HR = 1.91 with 95% CI 1.38 – 3.75; see Supplementary Fig. 6. Supplementary Fig. 7 shows that the resultant tumor subtypes were confounded by T-cell density, however, they demonstrated additional prognostic values with p = 0.003 (10×10 grid) and p = 0.004 (12×12 grid) after adjusting for T-cell density categorized into quartiles, based on multivariable Cox regression analyses.

In G-cross analysis, evaluating the likelihood of tumor cells having at least one T-cell within a specified radius (20 μm), similar T-cell density confounding effect was observed (Supplementary Fig. 8). Using a Cox regression model, we showed that tumors with the highest amount of T-cell infiltrates into both tumor epithelial and stromal areas i.e. Q4 showed significantly better survival than Q1 (univariable: p < 0.001; multivariable: p = 0.007), the significance was attenuated when it was restricted to T-cells found in the stromal region (univariable: p = 0.06; multivariable: p = 0.03); see Supplementary Fig. 9. Unlike MH, the prognostic significance of G-cross tumor subtypes was lost after adjusting for T-cell density categorized into quartiles, based on multivariable Cox regression analyses.

In contrast, the L-cross analysis, measuring the expected number of T-cells within a specified radius (20 μm) of a randomly chosen tumor cell, did not detect any survival significance based on the log-rank tests (Supplementary Fig. 10) as well as univariable and multivariable Cox regression analyses (Supplementary Fig. 11). All 3 estimators, namely isotropic, translation, and border, showed consistently insignificant prognoses. These less compelling survival results co-occurred with the absence of T-cell density correlation (Supplementary Fig. 10). Lastly, our data also showed that TIPC tumor subtypes had greater prognostic value as compared to that of T-cell density, MH index, G-cross, and L-cross functions (Table 1) based on multivariable Cox PH regression analyses.

**Table 1.** Prognostic significance of spatial subtypes (represented by clusters) identified by TIPC after adjusting for (a) CD3^+^ T-cell density, (b) Morisita-Horntumor:CD3^+^T cell index (10-by-10 pixels grid), (c) G-crosstumor:CD3^+^T cell AUC, and (d) L-crosstumor:CD3^+^T cell AUC, using multivariable Cox PH regression analyses. Tumor subtypes were grouped into quartile categories of (a) immune density, (b) MH index, (c-d) AUC of the G-cross and L-cross functions.

| Tumor subtypes |  | HR | lower 95% confidence interval | upper 95% confidence interval | p values |
| --- | --- | --- | --- | --- | --- |
| **CD3+ T-cell density** | 1st quartile [0,18] | Reference |  |  |  |
|  | 2<sup>nd</sup> quartile (18,77.8] | 1.09 | 0.80 | 1.49 | 0.593 |
|  | 3<sup>rd</sup> quartile (77.8,252] | 1.12 | 0.69 | 1.81 | 0.640 |
|  | 4<sup>th</sup> quartile (252,7.78e+03] | 1.51 | 0.75 | 3.04 | 0.252 |
| **TIPC** | cluster no.1 (S) | 0.95 | 0.70 | 1.30 | 0.770 |
|  | cluster no.2 (T) | Reference |  |  |  |
|  | cluster no.3 (TIL) | 0.56 | 0.33 | 0.94 | **0.029** |
|  | cluster no.4 (H&H) | 0.32 | 0.15 | 0.69 | **0.003** |
|  | cluster no.5 (SIL) | 0.71 | 0.37 | 1.36 | 0.301 |
|  | cluster no.6 (II-H) | 0.41 | 0.18 | 0.91 | **0.028** |

(b)
| Tumor subtypes |  | HR | lower 95% confidence interval | upper 95% confidence interval | p values |
| --- | --- | --- | --- | --- | --- |
| **Morisita-Horn index** | 1st quartile [0,0.07] | Reference |  |  |  |
|  | 2<sup>nd</sup> quartile (0.07,0.15] | 1.11 | 0.79 | 1.56 | 0.547 |
|  | 3<sup>rd</sup> quartile (0.15,0.25] | 1.18 | 0.81 | 1.72 | 0.399 |
|  | 4<sup>th</sup> quartile (0.25,0.80] | 0.79 | 0.48 | 1.31 | 0.362 |
| **TIPC** | cluster no.1 (S) | 1.02 | 0.73 | 1.42 | 0.904 |
|  | cluster no.2 (T) | Reference |  |  |  |
|  | cluster no.3 (TIL) | 0.68 | 0.43 | 1.06 | 0.085 |
|  | cluster no.4 (H&H) | 0.54 | 0.34 | 0.87 | **0.011** |
|  | cluster no.5 (SIL) | 0.87 | 0.54 | 1.41 | 0.577 |
|  | cluster no.6 (II-H) | 0.47 | 0.23 | 0.95 | **0.034** |

(c)
| Tumor subtypes |  | HR | lower 95% confidence interval | upper 95% confidence interval | p values |
| --- | --- | --- | --- | --- | --- |
| **G-cross** | 1st quartile [0,0.11] | Reference |  |  |  |
|  | 2<sup>nd</sup> quartile (0.11,0.44] | 0.98 | 0.71 | 1.35 | 0.892 |
|  | 3<sup>rd</sup> quartile (0.44,1.33] | 1.02 | 0.67 | 1.55 | 0.920 |
|  | 4<sup>th</sup> quartile (1.33,21.2] | 0.88 | 0.49 | 1.58 | 0.672 |
| **TIPC** | cluster no.1 (S) | 0.95 | 0.70 | 1.30 | 0.759 |
|  | cluster no.2 (T) | Reference |  |  |  |
|  | cluster no.3 (TIL) | 0.63 | 0.39 | 1.02 | 0.062 |
|  | cluster no.4 (H&H) | 0.51 | 0.28 | 0.92 | **0.026** |
|  | cluster no.5 (SIL) | 0.88 | 0.52 | 1.50 | 0.646 |
|  | cluster no.6 (II-H) | 0.46 | 0.22 | 0.96 | **0.039** |

(d)
| Tumor subtypes |  | HR | lower 95% confidence interval | upper 95% confidence interval | p values |
| --- | --- | --- | --- | --- | --- |
| **L-cross** | 1st quartile [0,339] | Reference |  |  |  |
|  | 2<sup>nd</sup> quartile (339,517] | 1.03 | 0.73 | 1.46 | 0.856 |
|  | 3<sup>rd</sup> quartile (517,669] | 0.91 | 0.64 | 1.29 | 0.587 |
|  | 4<sup>th</sup> quartile (669,1.58e+03] | 0.86 | 0.60 | 1.22 | 0.385 |
| **TIPC** | cluster no.1 (S) | 0.94 | 0.69 | 1.29 | 0.719 |
|  | cluster no.2 (T) | Reference |  |  |  |
|  | cluster no.3 (TIL) | 0.64 | 0.43 | 0.96 | **0.029** |
|  | cluster no.4 (H&H) | 0.47 | 0.31 | 0.69 | **<0.001** |
|  | cluster no.5 (SIL) | 0.84 | 0.53 | 1.34 | 0.465 |
|  | cluster no.6 (II-H) | 0.45 | 0.22 | 0.89 | **0.021** |

### Application of TIPC to characterize CD3^+^CD8^+^CD45RO^+^ T-cell spatial organization in TME

Leveraging the multiplexing capability of mIF assay, we applied TIPC to more specific memory cytotoxic T cells, an important anti-tumor phenotype, identified using three markers i.e. CD3^+^CD8^+^CD45RO^+^. Using *hex_len* = 80 pixels and *k* = 7 clusters (trend plots not shown), TIPC grouped the 930 tumors into 6 spatial subtypes with sizes ranging from 47 to 451, after excluding 1 tumor found in an outlier cluster. The 6 spatial subtypes are referred as: stroma-rich, tumor-rich, SIL-rich, TIL-rich, hot-and-homogeneous, and I:T high-I:S high (Fig. 6a, read from left to right), with cluster-median immune densities: 2.3, 1.2, 43.1, 43.6, 291.7, and 45.9 cells/mm^2^, accordingly (Fig. 6g).

**Figure 6.**
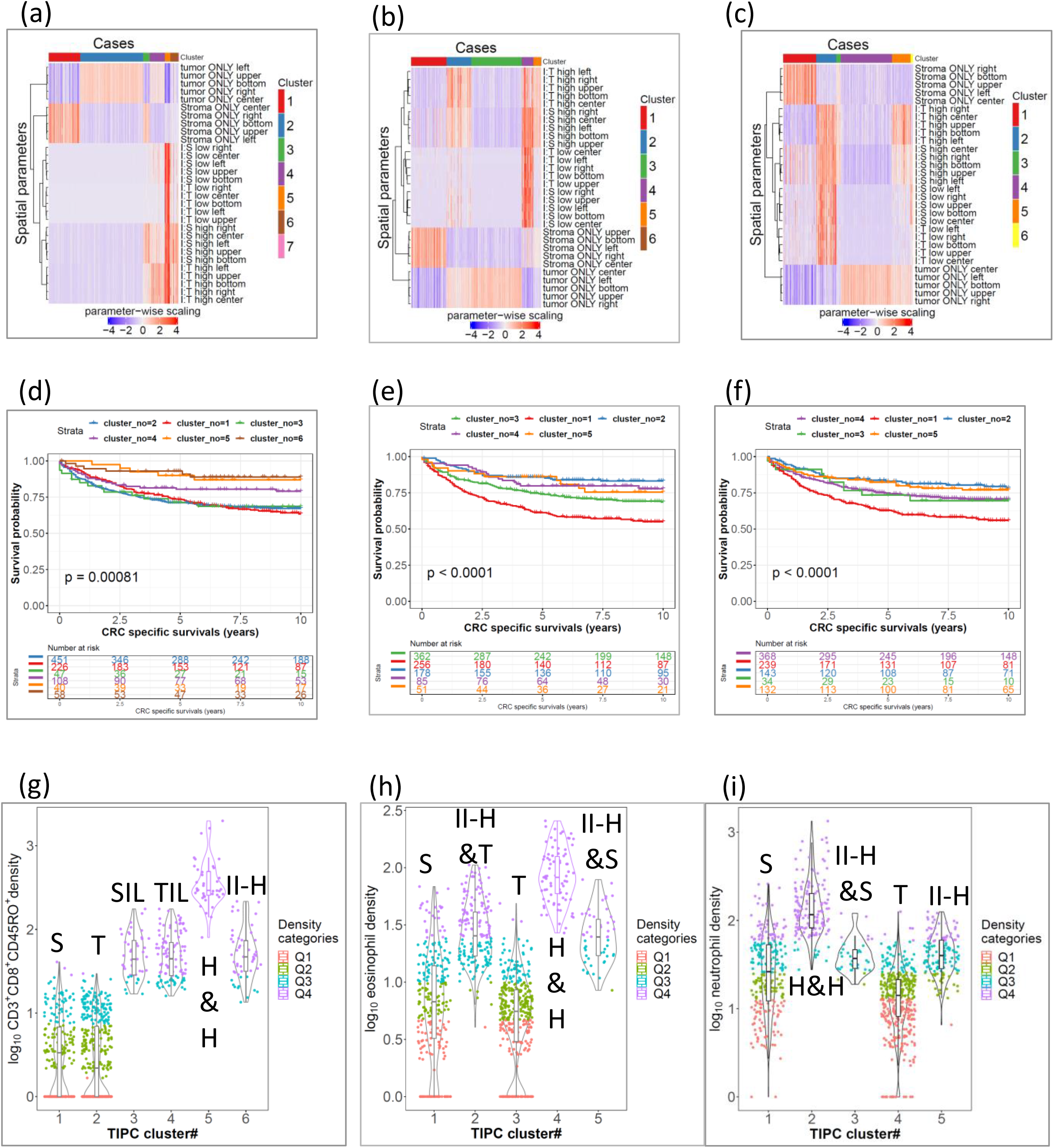
Application of TIPC to (**a**,**d**,**g**) CD3^+^CD8^+^CD45RO^+^ T-cell, (**b**,**e**,**h**) eosinophil, and (**c**,**f**,**i**) neutrophil CRC data. (**a**-**c**) Heat-maps display distinct spatial patterns, (**d**-**f**) Kaplan-Meier curves and log-rank test p-values demonstrate the prognostic relevance, and (**g**-**i**) boxplots (overlaid with violin plots) show overlapping and different immune density distribution across the resultant spatial subtypes where cases were color-coded by density quartile categories. In (**g**,**h**,**i**), jittered dots represent cases; box plot is defined by the 25th percentile (lower) and 75th percentile (upper) while extending lines mark the minimum (lower) and maximum densities (upper).

Our data showed that the resultant tumor subtypes were significantly associated with 10-year CRC-specific survival based on both the log rank test (p < 0.001; Fig. 6d) and Cox regression analyses (Supplementary Fig. 3). The univariable Cox regression analysis using tumor-rich as the reference revealed that TIL-rich, hot-and-homogeneous, and I:T high-I:S high exhibited better outcome. Using the multivariable Cox regression model to adjust for confounding factors, hot-and-homogeneous still showed significantly better CRC-specific survival (p = 0.04). Similar to the TIPC analysis of CD3^+^ T-cell data, the hot-and-homogeneous subtype (40 cases) exhibited 6.4 times higher CD3^+^CD8^+^CD45RO^+^ T-cell density than I:T high-I:S high subtype (58 cases) but these subtypes demonstrated similarly favorable clinical outcome as evinced by the overlapping Kaplan-Meier curves. We verified this assertion by conducting a chi-square test which showed insignificant differences between these two subtypes with 5-year CRC-specific survival (p > 0.05).

### Validation of TIPC using eosinophils identified from H&E-stained images

We then evaluated the extensibility of TIPC to characterize immune cells identified from H&E-stained tissue sections. Using eosinophils, with *hex_len* = 70 pixels and *k* = 6 clusters (trend plots not shown), TIPC grouped 934 tumors into 5 spatial subtypes with sizes ranging from 51 to 362, after excluding 2 tumors found in an outlier cluster. The spatial subtypes are referred as: stroma-rich, I:T high-I:S high-tumor-rich, tumor-rich, hot-and-homogeneous, and I:T high-I:S high-stroma-rich (Fig. 6b, read from left to right), with cluster-median eosinophil densities: 5.7, 24.4, 4.6, 81.4 and 23.8 cells/mm^2^, accordingly (Fig. 6h).

In spite of the substantially overlapping density distributions between hot-and-homogeneous and I:T high-I:S high, as well as between tumor-rich and stroma-rich, these subtypes were differentially associated with CRC survival (log rank p < 0.001, Fig. 6e). In the univariable Cox regression analysis, data showed that hot-and-homogeneous, I:T high-I:S high and even tumor-rich which had largely overlapping eosinophil density distribution exhibited significantly better CRC-specific survival (all p < 0.001 except for I:T high-I:S high with p = 0.01) than the reference stroma-rich. After adjusting for confounders using a multivariable Cox regression model, CRC-specific prognoses of hot-and-homogeneous (p < 0.001) and I:T high-I:S high-tumor-rich (p = 0.01) remained significant (Supplementary Fig. 3).

### Extended TIPC application to neutrophils identified from H&E-stained images

Utilizing the trend plots, tracking plots and heat-maps, an optimal TIPC solution for neutrophils was found at *hex_len* = 70 pixels and *k* = 6 clusters (Supplementary Fig. 12-14), as described below. Although *hex_len* = 70 pixels demonstrated slightly underrepresented I:T low component, it ensured minimal outlier tumors (i.e. 18) and also yielded slightly more effective clusters (5 clusters at *k* = 6), as compared to *hex_len* = 80 at its optimal *k* = 8 resulting in more than 40 outlier tumors; see Supplementary Fig. 14. Moreover, tumor subtypes determined using *hex_len* = 70, 80, and 100 pixels demonstrated largely similar spatial patterns (see Supplementary Fig. 14). As such, *hex_len* = 70 pixels and *k* = 6 were selected, where 916 tumors were grouped into 5 spatial subtypes with sizes ranging from 34 to 368, after excluding 18 tumors found in an outlier cluster. The tumor subtypes are referred as: stroma-rich, hot-and-homogeneous, I:T high-I:S high-stroma-rich, tumor-rich, and I:T high-I:S high (Fig. 6c, read from left to right), with corresponding cluster-median neutrophil densities: 25.2, 115.2, 36.1, 13.1, and 39.0 cell/mm^2^, accordingly (see Fig. 6i).

Kaplan-Meier curves (Fig. 6f) showed that stroma-rich represented the worst surviving tumor subtype, clearly separated from others (log rank p < 0.001). In the univariable Cox PH regression analysis, using stroma-rich as the reference, hot-and-homogeneous, tumor-rich, and I:T high-I:S high exhibited significantly better CRC-specific survival (all p values < 0.001). After adjusting for confounders using the multivariable Cox regression model, prognoses of hot-and-homogeneous (p = 0.004) and I:T high-I:S high (p = 0.01) remained significantly better than stroma-rich (Supplementary Fig. 3). Notably, tumor-rich, I:T high-I:S high and stroma-rich clusters exhibited substantially overlapping distributions (Fig. 6i). These results illustrated the ability of TIPC to distinguish amongst tumors even with similar immune densities.

### Consistent TIPC results over a range of parameter settings

TIPC demonstrated robust performance over a range of sub-region sizes (*hen_len* = 60 to 110 pixels equivalent to 2301 to 7733 square micrometers at 20x objective image scanning) and number of clusters (*k* = 4 to 10). Results are shown as follows: Fig. 5b and Supplementary Fig. 5 for CD3^+^ T-cell analysis, and Supplementary Fig. 15 for neutrophils analysis, which summarized Cox regression analysis results using tumor subtype of the lowest median immune density as the reference. Our data showed that TIPC was able to produce tumor subtypes with similar sample sizes and spatial patterns, and, importantly, these tumor subtypes showed consistent prognostic association. In CD3^+^ T-cell analysis, the hot-and-homogeneous subtype showed consistently better prognosis than the reference which was either tumor-rich or stroma-rich (as they had similarly low immune densities). Additionally, both TIL-rich and I:T high-I:S high subtypes also showed more favorable CRC survival than the reference most of the time though the former was more robust than the latter. In neutrophil analysis, stroma-rich tumors showed consistently worse prognosis than the tumor-rich tumors despite having substantially overlapping immune densities.

## Discussion

Recent advancement of multiplexed tissue imaging techniques empowers deep phenotyping of immune cells, along with their spatial locations in the TME, it generates a valuable resource for advancing the understanding of tumor immunity^14^. To address the lacking of effective data analysis methods, we developed a novel computational algorithm, named as Tumor-Immune Partitioning and Clustering (TIPC), for characterizing tumor-immune spatial relationship in TME.

Existing studies have been mainly focusing on assessing the prognostic significance of immune cell density, while offering direct clinical translation, it underutilized the valuable spatial information^18, 19^. Our data showed that TIPC was capable of identifying tumor subtypes with unique tumor-immune spatial patterns, more importantly, these tumor subtypes demonstrated prognostic value which were otherwise undetectable using immune density data alone. TIPC applications to eosinophil and neutrophil revealed two immune-depleted subtypes with similarly low immune densities, termed tumor-rich and stroma-rich, demonstrating differential CRC survival outcomes. These results illustrated the ability of TIPC in capturing not only immune cell spatial distribution, but also tumor morphology which can have important clinical and biologic implications.

Although increasing efforts have been undertaken to go beyond immune densities, using CD3^+^ T-cell data, we showed that Morisita-Horn index and G-cross function were largely confounded by the immune density whereas L-cross function did not show prognostic significance. After adjustment for immune density, only MH-based tumor subtypes remained significantly associated with CRC-specific survival but not G-cross. Masugi et al. also pointed out that L-cross function is limited by the underlying assumption that tumor cells are treated as isolated entities, without considering tumor morphology like ductular structures in pancreatic adenocarcinoma^12, 20^. To capture the highly heterogeneous tumor-immune cell interactions within a TME, TIPC employs novel spatial measures quantified on tessellated sub-regions. As such, TIPC yields tumor subtypes encapsulating intrinsic information of immune cell densities, tumor-immune spatial relationship, and tumor morphology. We showed that TIPC tumor subtypes provided additional prognostic value to immune density, and also outperformed existing methods.

To our knowledge this is the first study examining the association between spatially informed tumor subtypes and histologic and molecular features of cancer using large-scale cohort data. Using CD3^+^ T cells, TIPC found tumor subtypes with significant association with well-established CRC prognosticators including AJCC disease stage, lymphocytic reactions, MSI status, and CIMP status, providing corroborative evidence for its clinical relevance. Moreover, TIPC was able to reveal heterogeneous T-cell response within MSI-high phenotype and demonstrated a strong association with immune checkpoint markers i.e. PDCD1 and PDCD1LG2. Taken together, TIPC can potentially be used to inform patient selection for immune checkpoint therapy.

Importantly, TIPC performance was robust to the choice of immune cell types, detection methods and parameter settings. Besides CD3^+^ T-cell, using a more specific CD3^+^CD8^+^CD45RO^+^ T-cell subset^21, 22^ and two innate immune cell types i.e. eosinophils and neutrophils, TIPC was still able to determine prognostic spatial subtypes. With eosinophils and neutrophils, we showed that TIPC is extensible to immune cells identified from routinely H&E stained images. We also tested the robustness of TIPC performance over a range of sub-region sizes and different number of clusters, using both CD3^+^ T-cell and neutrophil data. TIPC successfully identified tumor subtypes exhibiting similar spatial patterns and consistent prognostic significance across the tested parameter values.

There were several limitations in our study. First, due to the unavailability of treatment information, its potentially confounding effect was not accounted for in the prognostic analyses. Second, like other unsupervised clustering algorithms, the consensus hierarchical clustering adopted in TIPC requires a reasonable sample size for the identification of robust spatial subtypes with sufficient statistical power for downstream analysis. The sample size is both data-dependent (for instance cancer type) and study design-dependent (for instance hypothesis testing versus exploratory analysis). Third, TIPC is limited to the study of single immune cell type at a time. Fourth, unlike histoCAT ^23^ taking raw mass cytometry images as input, TIPC requires input data containing both the identity and location of tumor, stromal, and immune cells, for two major reasons: (1) the accessibility to well-established cell segmentation and phenotyping algorithms like InForm, QuPath, and CellProfiler is convenient, and (2) it prevents rigid input data format. Fifth, the TMAs were only limited to cores from the tumor center, impeding assessment of the prognostic association of immune cell infiltrate at the invasive margin ^24^.

In conclusion, our proposed TIPC algorithm provides an effective computational solution for identifying tumor subtypes with unique tumor-immune spatial patterns. The resultant spatial subtypes uniquely and concisely encapsulate: (1) the relative distribution between immune and tumor (or stromal) cells (i.e. partitioning); (2) the proximity among immune cells (i.e. clustering); (3) tumor morphology, for instance, well differentiated tumors tend to have a relatively higher proportion of tumor-only sub-regions. While immune density is not considered explicitly, it is also captured by TIPC thus supplementing extra information. For instance, tumors enriched in I:T high or I:S high sub-regions tend to have more immune cells than tumors enriched in tumor-ONLY or stroma-ONLY sub-regions. Through identifying spatially-informed tumor subtypes, TIPC fosters a deeper understanding of the interplay of multiple tumor and microenvironmental features in cancer immunology. Given the easy-to-use TIPC R package freely available online, TIPC implementation can be easily incorporated into pre-clinical and clinical studies, serving as a tumor-immune spatial profiling tool for quantifying changes in immune contexture at different stages of tumorigenesis and in response to different treatments.

## Online Methods

### Patient selection

We have been collecting data from two prospective cohort studies in the U.S., the Nurses’ Health Study (NHS, 121,701 women aged 30-55 years followed since 1976) and the Health Professionals Follow-up Study (HPFS, 51,529 men aged 40-75 years followed since 1986). ^25^ Every two years, study participants have been followed with questionnaires to collect information on lifestyle factors and medical history including colorectal cancer. The National Death Index was used to ascertain deaths of study participants and identify unreported lethal colorectal cancer cases. Participating physicians reviewed medical records to confirm diagnosis of colorectal cancer, and to record data on tumor characteristics including tumor anatomical location, disease stage based on the American Joint Committee on Cancer TNM classification. Formalin-fixed paraffin-embedded (FFPE) tissue blocks were collected from hospitals where participants diagnosed with colorectal cancer had undergone tumor resection. We included 931 patients with available tissue microarray (TMA) of colorectal carcinoma tissue diagnosed up to 2008. Our TMAs included up to four cores from colorectal cancer and up to two cores from normal tissue blocks, as described previously.^26^ We included both colon and rectal carcinomas, on the basis of the colorectal continuum model.^27^ A single pathologist (S.O.), who was unaware of other data, conducted a centralized review of hematoxylin and eosin-stained tissue sections of all colorectal carcinoma cases, and recorded pathological features including tumor differentiation and lymphocytic reaction patterns [tumor-infiltrating lymphocytes (TIL), intratumoral periglandular reaction, peritumoral lymphocytic reaction, and Crohn’s-like lymphoid reaction].^28^ Tumor differentiation was categorized as well to moderate or poor (> 50% vs. ≤ 50% glandular area, respectively). As previously described, each of four lymphocytic reaction components was graded as absent/low, intermediate, or high.^28^ Informed consent was obtained from all participants at enrolment. This study was approved by the institutional review boards at Harvard T.H. Chan School of Public Health and Brigham and Women’s Hospital (Boston, MA).

### Immunohistochemical analyses

Immunohistochemical analyses of CDH1 (E-cadherin), nuclear CTNNB1 (beta-catenin), PDCD1 (programmed cell death 1, PD-1), and PDCD1LG2 (programmed death 1 ligand 2, PD-L2), PTGER2 (prostaglandin E receptor 2), SQSTM1 (p62), and YAP1 were performed using anti-CDH1 antibody (Dako, Carpinteria, CA), anti-CTNNB1 antibody (BD Transduction Laboratories, Franklin Lakes, NJ), anti-PDCD1 antibody (clone EH33), anti-PDCD1LG2 antibody (clone 366C.9E5), anti-PTGER2 (Cayman Chemical, Ann Arbor, MI), anti-SQSTM1 (Abnova, Taipei, Taiwan;), and anti-YAP1 antibody (Cell Signaling, Danvers, MA), respectively.^29–34^ Anti-PDCD1 antibody and anti-PDCD1LG2 antibody were generated in the laboratory of G.J. Freeman at Dana-Farber Cancer Institute.^28^

### Evaluation of Tumor Molecular Characteristics

Genomic DNA was extracted from colorectal carcinoma tissue in whole-tissue sections of archival FFPE tissue blocks using QIAamp DNA FFPE Tissue Kit (Qiagen, Hilden, Germany). MSI status was determined using polymerase chain reaction (PCR) of 10 microsatellite markers (D2S123, D5S346, D17S250, BAT25, BAT26, BAT40, D18S55, D18S56, D18S67, and D18S487), and MSI-high was defined as presence of instability in ≥ 30% of the markers, as previously described.^26^ Methylation status of eight CpG island methylator phenotype (CIMP)-specific promoters (*CACNA1G*, *CDKN2A*, *CRABP1*, *IGF2*, *MLH1*, *NEUROG1*, *RUNX3*, and *SOCS1*) and long-interspersed nucleotide element-1 (LINE-1) were determined using MethyLight assay on bisulfite-treated DNA.^27^ CIMP-high was defined as ≥ 6 methylated promoters of eight promoters, and CIMP-low/negative as 0-5 methylated promoters, as previously described.^27^ PCR and pyrosequencing were performed for *KRAS* (codons 12, 13, 61, and 146), *BRAF* (codon 600), and *PIK3CA* (exons 9 and 20), as previously described.^27^ Whole exome sequencing was performed using DNA from tumor and matched normal tissue pairs, as previously described.^35^ Using a neoantigen prediction pipeline for somatic mutations, the neoantigen load (i.e., the number of proteins that likely give rise to immunogenic peptides in the tumor microenvironment) was estimated by counting peptides that bind to personal HLA (human leukocyte antigen) molecules with high affinity (< 500 nM). Using NetMHCpan (version 2.4),^36^ we predicted the binding affinities of all possible 9- and 10-mer mutant peptides to the corresponding HLA alleles inferred by the POLYSOLVER algorithm, as previously described.^35^

### T-cell multiplex immunofluorescence staining

As mentioned above, we constructed tissue microarrays of colorectal cancer cases with sufficient tissue materials, including up to four tumor cores from each case in one tissue microarray block. Deparaffinized 4 μm sections from TMA blocks were incubated with the primary antibody, then treated with anti-mouse/rabbit horseradish peroxidase-conjugated (HRP) secondary antibody (Opal Polymer; PerkinElmer, Hopkinton, MA), and finally incubated with the fluorophore amplification reagent (PerkinElmer) for tyramide signal amplification. The slides were sequentially stained using the following antibodies/fluorescent dyes, in order: anti-CD3 antibody (clone F7.2.38; Dako; Agilent Technologies, Carpenteria, CA)/Opal-520, anti-FOXP3 (clone 206D, Biolegend, San Diego, CA)/Opal-540, anti-CD45RO (clone UCHL1, Dako)/Opal-650, anti-CD8 (clone C8/144B, Dako)/Opal-570, anti-CD4 (clone 4B12, Dako)/Opal-690, anti-KRT (Keratins, pan-keratins)(clone AE1/AE3, Dako) in combination with anti-KRT (clone C11, Cell signaling, Denvers, MA)/Opal-620. Each slide was then treated with spectral nuclear counterstain 14’,6-diamidino-2-phenylindole (DAPI) (clone FP1490, PerkinElmer). Fully stained slides were stored in a lightproof box at 4°C prior to imaging. T-cell multiplex immunofluorescence image acquisition and analysis – tissue and cell segmentation, and cell phenotyping

Digital images of all TMA cores were acquired using Vectra 3.0 quantitative pathology imaging system (PerkinElmer) equipped with 20× objective. Large areas with necrosis, artefact, or excessive tissue folding were excluded from analysis. Demultiplexed images of each core underwent first tissue segmentation to characterize regions of tumor epithelium and peritumoral stroma based on cytokeratin expression, using supervised machine learning algorithms within Inform 2.4.1 (PerkinElmer). Following tissue segmentation, cell enumeration and segmentation was performed using the DAPI signal to aid in identification of nuclei. Each cell was further segmented into nuclear, cytoplasmic and membranous compartments. A separate supervised machine learning algorithm was used to identify T-cells based upon a combination of cytomorphology and subcellular T-cell marker expression patterns. These single cell data were then used to calculate T-cell subpopulation densities within separate regions. Aggregate tumor-level densities were then determined by calculating the average density for each T-cell subset across all cores from each tumor.

### Hematoxylin and eosin based eosinophil and neutrophil phenotyping

We used QuPath, an open source software platform for whole slide image analysis, to identify tumor cells, eosinophils and neutrophils in images of H&E stained colorectal cancer tissue microarray sections.^37^ The H&E stained tissue microarray sections were scanned using Vectra 3.0 quantitative pathology imaging system (PerkinElmer) equipped with 20× objective. The image analysis steps were based on basic functions of QuPath: (1) *estimate stain vectors* to extract the H&E stain vectors and background values from the images; (2) *simple tissue detection* to discriminate the tissue region from the white background; (3) *cell detection* to detect cells based on the size, shape, and optical density of nuclei in the hematoxylin layer and to calculate features of the cells including nuclear area and circularity; (4) *add smoothed features* to calculate Gaussian-weighted mean of the cell measurements in the neighboring cells; and (5) *create detection classifier* to train a random forests classifier using 150 image subset to identify the cell types of interest. These commands were recorded as a script, which was run across all the images.

### Tumor-Immune Partitioning and Clustering (TIPC) analytical approach

Tumor-Immune Partitioning and Clustering (TIPC) is a novel computational method that utilizes hexagonal tessellation and a classifier that evaluates multiple spatial parameters against a tumor region-specific null model representing a state of neutral tumor-immune cell interactions. Fig. 2 dictates the major analysis components of TIPC using an example of multiplexed IF stained image. The TIPC R package is freely available on the web https://github.com/MPE-Lab.

In a two-dimensional (2D) tissue section space where the identity and location of tumor, non-tumor (i.e. stromal), and immune cells are known, TIPC first divides the space into hexagonal sub-regions (spatstat R package^38^) and calculates two global ratios using the total number of immune, tumor and stromal cells representing a state of neutral tumor-immune cell interactions. The sub-regions are then classified into 6 different categories, namely tumor-only, immune-to-tumor low (I:T low), immune-to-tumor high (I:T high), stroma-only, immune-to-stroma low (I:S low), and immune-to-stroma high (I:S high), based on the comparing of immune, tumor and stromal cell content of each sub-region to the global immune to tumor and immune to stroma ratios. For example, if a sub-region contains only tumor cells, it will be categorized as tumor-only, whereas if there are immune cells found in the same sub-region with tumor cells, but the local immune to tumor ratio is smaller (or larger) than the global ratio, the sub-region will be categorized under the I:T low (or I:T high) category. The three stromal categories are defined in a similar way. See Supplementary Table 2 for the summary of definitions. After counting the number of sub-regions in each category, the counts are normalized using the total number of non-empty or informative sub-regions. As such, the resultant six-element vector (hereafter called TIPC spatial parameters) captures the essence of tumor-immune spatial pattern of the tumor microenvironment, and it can be readily compared across tumors.

To identify tumor subtypes with similar spatial patterns (abbreviated as spatial subtypes), TIPC employs a consensus hierarchical clustering algorithm to group tumors in an unsupervised manner, where the input data is a matrix formed by concatenating the six-element spatial parameters of all the tumors in the entire cohort. Based upon the ConsensusClusterPlus R package,^39^ the *consensus_clustering* function in TIPC R package allows users to adjust the following parameters, (1) distance function: Pearson correlation (default); (3) number of clusters *k*: 2 to 6 (default); (4) number of repeated clustering to reach consensus: 50 (default); (5) proportions of cases i.e. tumors to be included in each repeat: randomly selected 80% (default); (6) proportions of features to be included in each repeat: all six spatial parameters (default). At a specified *k*, clustering is repeated for 50 times and a cumulative consensus measure matrix is determined by computing the number of times each tumor is assigned to the same cluster with each of all other tumors. Final cluster assignment is determined through applying a hierarchical agglomerative clustering with complete linkage on the cumulative consensus matrix. Two main reasons of adopting hierarchical clustering than other approaches, such as k-means, are the companion heat-map (1) provides a visual inspection of the tumor clustering quality i.e. cohesion spatial patterns within the same cluster while distinct spatial patterns between clusters, and it (2) facilitates the interpretation of resultant tumor-immune spatial patterns. See Supplementary Table 3 for the description of a list of characteristic spatial subtypes identified by TIPC in this study.

To facilitate the determination of optimal number of clusters (*k*), TIPC R package inherits several auxiliary functions and plots from the ConsensusClusterPlus R package. The consensus cumulative distribution function (CDF) delta curve displays the amount of additional consensus being gained at every increment of *k*. In a CDF plot, the minimum number of stable clusters i.e. spatial subtypes can be found at the location where the gain starts to diminish. The tracking plot further enables examination of cluster stability by tracing the cluster assignment of individual tumors as the number of clusters (*k*) increases. Lastly, heat-map of the six spatial measures with tumors ordered by their subtype assignment serve as a visual confirmation of clustering quality. An optimal choice of *k* is data-dependent, for instance, tumor types and immune cell of interest.

Some tumors might be assigned to isolated and small clusters (hereafter called outlier clusters) as a result of suboptimal choice of parameter values or rare spatial patterns which can be biologically true or technical artefacts. Even supposing it is a true spatial pattern, an outlier cluster contains very few tumors and hence losing its statistical power in downstream analysis. To facilitate determining of optimal *k* and *hex_len* that minimize the number of outlier tumors, TIPC R package provides two trend plots: (1) one shows the total number of outlier tumors, and (2) another shows the number of outlier clusters alongside with the number of effective clusters, as a function of *k* and *hex_len*. The Cox proportional hazards (PH) regression analysis function, *postTIPC_SurvivalAnalysis*, excludes clusters contain less than 30 tumors by default.

As a rule of thumb, an optimal sub-region size *hex_len* should ensure tumor-immune spatial relationship represented by the six spatial parameters are well captured i.e. not biased towards specific parameters. To examine the effect of *hex_len*, TIPC R package incorporates three auxiliary functions, namely *multiple_hexLen_tessellation*, *multiple_hexLen_count_TIPC_cat*, and *trend_plot_hexLen*. The former two are wrapper functions: (1) *multiple_hexLen_tessellation* for dividing the 2D space into sub-regions, and (2) *multiple_hexLen_count_TIPC_cat* for counting the number of sub-regions in each of the six categories, across a specified range of *hex_len*. *trend_plot_hexLen* consolidates these data and produces a trend plot showing relative composition of the six sub-region categories, as a function of *hex_len*.

To minimize the sensitivity to sub-region size selection, we proposed a grid (i.e. a collection of sub-regions) shifting approach aiming to capture extended scale of information without needing an excessively large sub-region size. Here, we refer to the entire set of sub-regions formed on a 2D image space as a grid. At a specified *hex_len*, five sets of the six-element TIPC spatial parameters are computed using five different grids generated as follows. A central grid is created by populating from the center location of the 2D space (spatstat R package *hextess* function with default zero offset values in both XY locations), and the other four grids are created by shifting to the left, right, upper, and lower directions at a distance equals to half of the *hex_len* away from the central grid. A trend plot for visual inspection of the effect of grid shifting on the TIPC spatial measures is available in the R package by calling the function *trend_plot_shiftDirection*.

### Statistics and computational algorithms

All analyses were conducted using statistical software R (version 3.6.1). We used the two-sided α level of 0.05. We evaluated the relationship of TIPC clusters (alternatively also referred as tumor subtypes or spatial subtypes) with 10-year colorectal cancer-specific survival outcome using both univariable and multivariable Cox proportional hazards (PH) regression models (survival R package). The cluster assignment was taken as a categorical variable and hazard ratio (HR) of each cluster against the reference cluster was estimated from the Cox PH regression models. In the robustness analysis, cluster with the lowest immune density was used as the reference. To assess the overall survival significance of spatial subtypes, we employed Kaplan-Meier method to estimate cumulative survival probabilities which were then compared statistically using the log-rank test (survminer R package). Note that in these analyses of colorectal cancer-specific mortality, deaths resulting from other causes were censored.

In multivariable Cox PH regression analyses, covariates assessed as potential confounding factors included sex (female vs. male), age at diagnosis (continuous), year of diagnosis (continuous), family history of colorectal cancer in any first-degree relative (present vs. absent), tumor location (proximal colon vs. distal colon vs. rectum), tumor differentiation (well to moderate vs. poor), disease stage (I/II vs. III/IV), MSI status (MSI-high vs. non-MSI-high), CIMP status (high vs. low/negative), LINE-1 methylation level (continuous), *KRAS* mutation (mutant vs. wild-type), *BRAF* mutation (mutant vs. wild-type), and *PIK3CA* mutation (mutant vs. wild-type). Cases with missing data were included in the majority category of a given categorical covariate to limit the degrees of freedom: family history of colorectal cancer in a first-degree relative (0.4%), tumor location (0.4%), tumor differentiation (0.1%), disease stage (7.2%), MSI (2.9%), CIMP (7.1%), *KRAS* (2.9%), *BRAF* (2.0%), and *PIK3CA* (8.7%). For the cases with missing LINE-1 methylation data (2.8%), we assigned a separate indicator variable. We performed feature selection for covariates by fitting a generalized linear model (using Cox regression model as the link function) with lasso regularization (glmnet R package), whereby the optimal regularization parameter lambda was selected from a 5-fold cross validation test. Our data showed that similar p-values, HRs, and CIs were obtained for the cluster variable after the covariate selection (data not shown).

We employed the Cochran-Armitage trend test^40, 41^ to examine for the association relationship between spatial subtypes (categorical variable) and the clinicopathologic and molecular features (ordinal variable) which include 5-year CRC-specific survival, microsatellite instability (MSI) status, CpG island methylator phenotype (CIMP) status, lymphocytic reactions^28^, neoantigen load^35^, and immunohistochemistry (IHC) based protein expression of CDH1, CTNNB1, PDCD1, PDCD1LG2, PTGER2, SQSTM1, and YAP1.

We compared the performance of TIPC with state-of-the-art analysis methods, including multi-type G-function (G-cross), multi-type L-function (L-cross), and Morisita-Horn (MH) index, using CD3^+^ T-cell data. The former two methods are used as a surrogate to quantify relative proximity between two selected cell types, in this case, tumor cells and CD3^+^ T-cells. The latter method allows us to directly quantify the degree of co-localization of tumor cells and CD3^+^ T-cells. Considering that proximity to tumor cells is of less importance once T-cells already infiltrate into tumor epithelial area, we conducted these analyses using (1) all T cells, and also (2) restricted to T cells detected in tumor stromal region.

In MH analysis, we first calculated the MH indices (spaa R package) of individual cases using four rectangular grids of sizes 9-by-9, 10-by-10, 11-by-11, and 12-by-12 (in pixels) (spatstat R package). Adopting the approach proposed in the original work^15^, the entire cohort was randomly divided into two equal portions for discovery and validation purpose. Subsequently, 13 cut-off values equally spaced from 20^th^ to 80^th^ percentiles using 5^th^ percentile step size were tested, whereby the cases in the discovery subset were dichotomized into MH-high and low groups. At each of these cut-off values, a univariable Cox PH regression analysis was conducted to assess if the MH-high tumor subtype was prognostically different from the MH-low tumors within the discovery subset. The cut-off values generated from discovery subset was then tested in the validation subset. Benjamini and Hochberg method was employed to control for false discovery rate (FDR) resulting from multiple comparisons of the 13 cut-off values. Of note, cases with no detection of CD3^+^ T-cell were excluded in the survival analysis.

G-cross^16^ and L-cross^12^ functions are generalizations of the G- and L-functions, which are nearest neighbor distance functions, to multi-type point patterns. These functions are based on the theory of point process. In this study, we used G-cross to estimate the probability of finding at least one CD3^+^ T-cell *j* within a *r* radius of any tumor cell *i*. On the other hand, L-cross function is a transformed Ripley’s K-function taking the form of 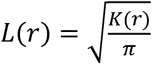. Such transformation provides two major advantages, firstly the square root stabilizes the variance of the estimator, and secondly it simplifies interpretation as the theoretical value of L(*r*) = *r* in case of a completely random (uniform Poisson) point pattern. The multi-type K-function, which underlie the L-cross function, is defined as 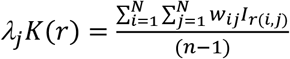 where the right side of the equation represents the expected number of additional CD3^+^ T-cells *j* within a distance *r* of a randomly chosen tumor cell *i*, and *w_ij_* is the edge correction weight. The estimation of G-, K-, and L-function is impeded by edge effects due to the unobservable points outside the data window, hence, an edge correction is required. There are several edge correction methods incorporated in the spatstat R package. In this study, we applied Kaplan-Meier (km) method for G-cross estimation as the other two estimators, i.e. reduced sample (rs) and Hanisch-style (han), failed to model some of the cases (data not shown). As for L-cross analysis, we tested all the available options including isotropic, translation, and border estimators. To quantify the degree of CD3^+^ T-cell infiltration, the area under the curve (AUC) of G-cross and L-cross functions were computed from 0 to *r* = 20 micrometers (equivalent to 0 to 40 pixels at 20x objective image scanning) considering that 20 micrometers are the longest intercellular distance for effective CD3^+^ T-cell to tumor cell interactions^16^. For downstream survival analysis, patients were grouped into ordinal AUC quartile categories i.e. Q1 to Q4 from low to high AUC values.

## Acknowledgements

Portions of this research were conducted on the O2 High Performance Compute Cluster, supported by the Research Computing Group, at Harvard Medical School (see http://rc.hms.harvard.edu for more information).

We would like to thank the participants and staff of the Nurses’ Health Study and the Health Professionals Follow-up Study for their valuable contributions as well as the following state cancer registries for their help: AL, AZ, AR, CA, CO, CT, DE, FL, GA, ID, IL, IN, IA, KY, LA, ME, MD, MA, MI, NE, NH, NJ, NY, NC, ND, OH, OK, OR, PA, RI, SC, TN, TX, VA, WA, WY. The authors assume full responsibility for analyses and interpretation of these data.

## Author Contributions

M.C.L., S.O. and J.A.N. developed the main concept and designed the study. M.C.L., J.P.V., J.B. and J.A.N. designed the computational algorithm. C.S.F., S.O., and J.A.N. wrote grant applications. J.B., J.P.V., K.H., A.D.C., A.D.S., K.A., T.H., R.N., S.O., and J.A.N. were responsible for collection of tumour tissue, and acquisition of epidemiologic, clinical and tumour tissue data, including histopathological and immunohistochemical characteristics. M.C.L., J.P.V., M.Z., S.G., and K.H. contributed to data analysis and interpretation. M.C.L., J.B., J.P.V., K.H., S.O., and J.A.N. drafted the manuscript. S.O. and J.A.N. supervised the project. K.F., R.N., J.K.L., C.S.F., C.J.W., S.O., and J.A.N. contributed to editing and critical revision for important intellectual content. All authors reviewed and approved the manuscript.

## Use of Standardized official symbols

We use HUGO (Human Genome Organisation) Gene Nomenclature Committee (HGNC)-approved official symbols (or root symbols) for genes and gene products, including BRAF, CACNA1G, CD3, CD4, CD8, CDKN2A, CDH1, CRABP1, CTNNB1, FOXP3, IGF2, KRAS, KRT, MLH1, NEUROG1, PDCD1, PDCD1LG2, PIK3CA, PTGER2, PTPRC, RUNX3, SOCS1, SQSTM1, and YAP1; all of which are described at www.genenames.org. Gene symbols are italicized whereas symbols for gene products are not italicized.

