## Supplementary figures and tables for "Tumor-Immune Partitioning and Clustering (TIPC) algorithm reveals distinct signatures of tumor-immune cell interactions within the tumor microenvironment"

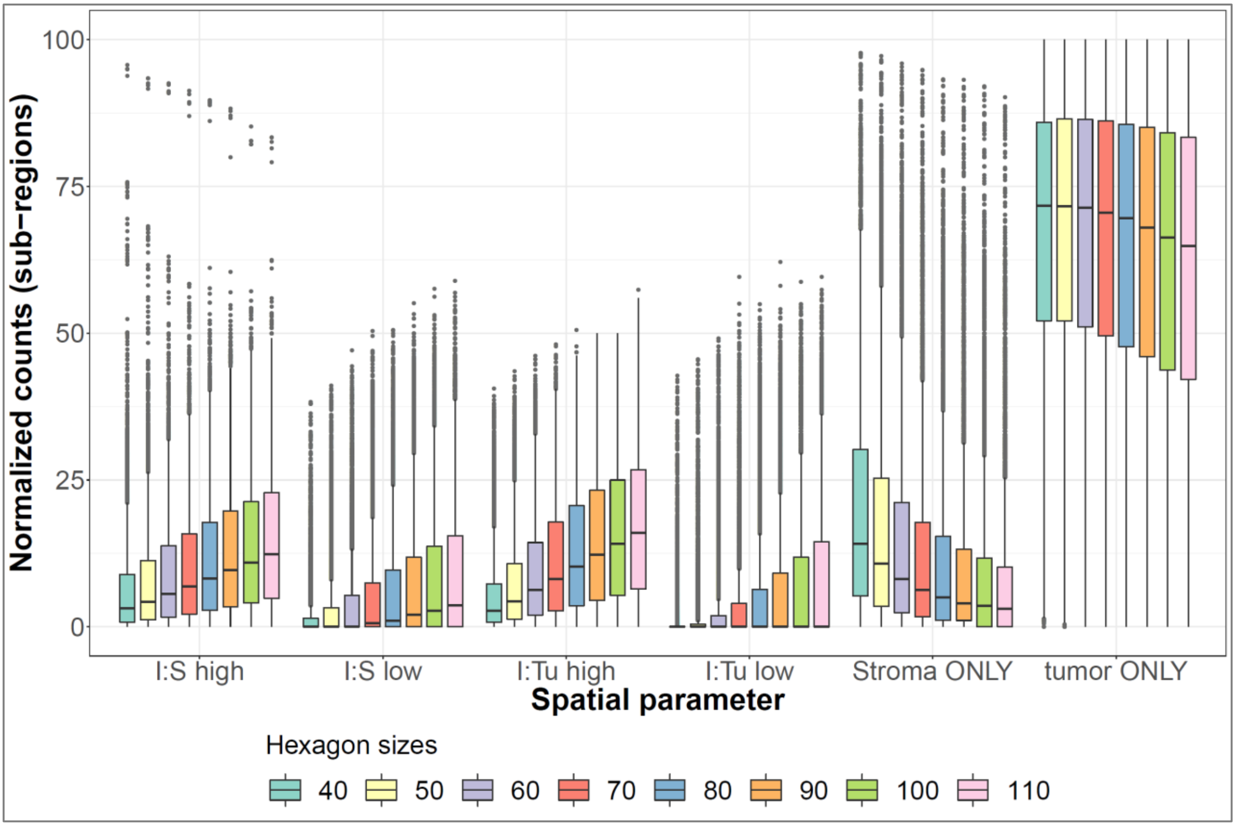


**Supplementary Figure 1.** Trend plot shows the distribution of %composition of the six TIPC spatial parameters at different sub-region sizes (i.e. *hex_len*) based on CD3$^{+}$ T-cell CRC data. Box plot is defined by the 25th percentile (lower) and 75th percentile (upper) while extending lines mark the minimum (lower) and maximum densities (upper); gray dots represent cases.

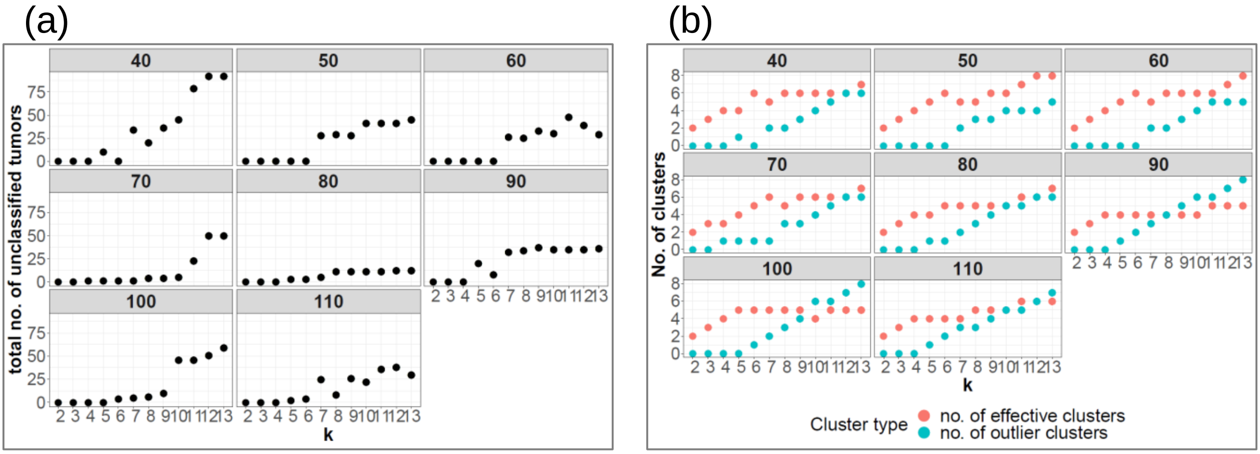


**Supplementary Figure 2.** Trend plots show (**a**) the total number of unclassified tumors i.e. those were detected in outlier clusters, (**b**) the number of outlier and effective (i.e. major clusters) clusters, obtained at different sub-region sizes (*hex_len* = 40 to 110, at interval of 10 pixels) and number of clusters (*k*). TIPC analysis was conducted using $CD3^+$ T-cell CRC data. Outlier clusters were defined as clusters comprised of fewer than 30 tumors.

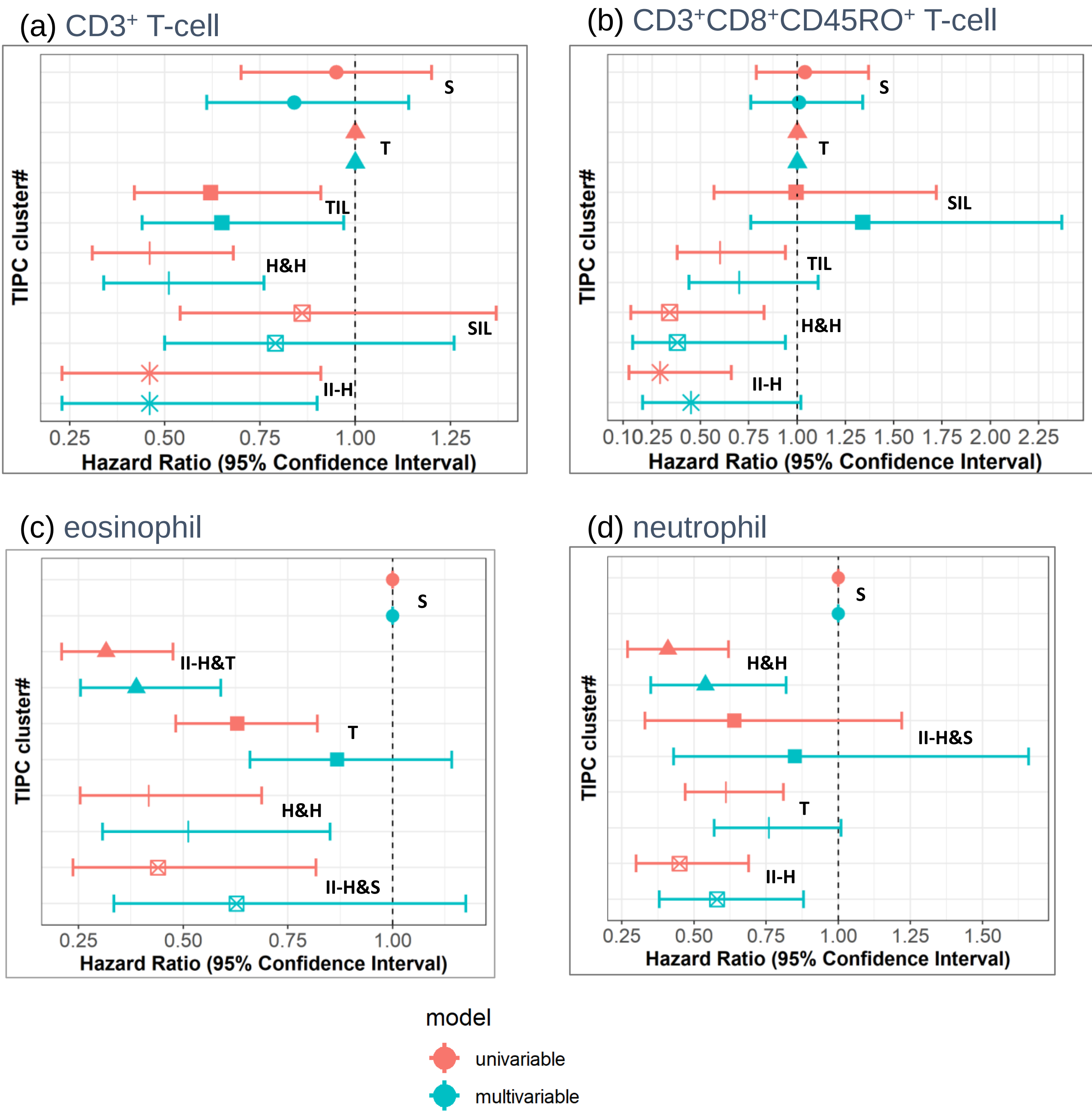


**Supplementary Figure 3.** Forest plots of spatial subtypes determined by TIPC using (**a**) $CD3^+$ T-cell CRC data at *hex_len* = 70 and *k* = 9, (**b**) $CD3^+CD8^+CD45RO^+$ T-cell at *hex_len* = 80 and *k* = 7, (**c**) eosinophil at *hex_len* = 70 and *k* = 6, (**d**) neutrophil at *hex_len* = 70 and *k* = 6, using cox proportional hazards analyses. Characteristic spatial subtypes: S = stroma-rich, T = tumor-rich, TIL = TIL-rich, II-H = I:T high-I:S high, H&H = hot-and-homogeneous, II-H&T = I:T high-I:S high-tumor-rich, and II-H&S = I:T high-I:S high-stroma-rich. The thin horizontal lines i.e. whiskers indicate the magnitude of the confidence interval: lower 95% (left) and upper 95% (right).

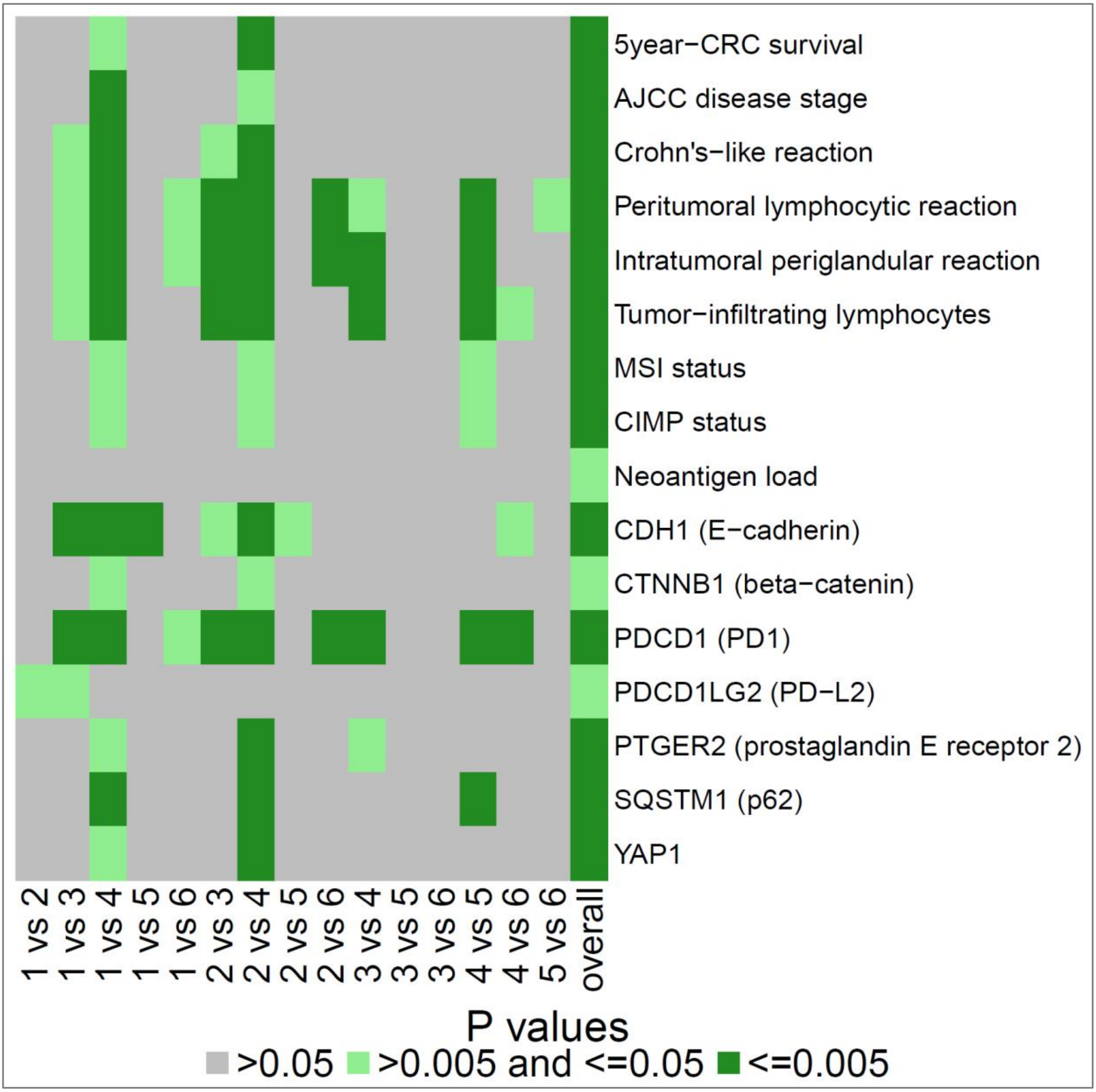


**Supplementary Figure 4.** Extended Cochran–Armitage test of association for TIPC clusters (unordered variable) and clinicopathological and molecular features (ordered variable). TIPC analysis was performed using $CD3^+$ T-cell CRC data at *hex_len* = 70 and *k* = 9.

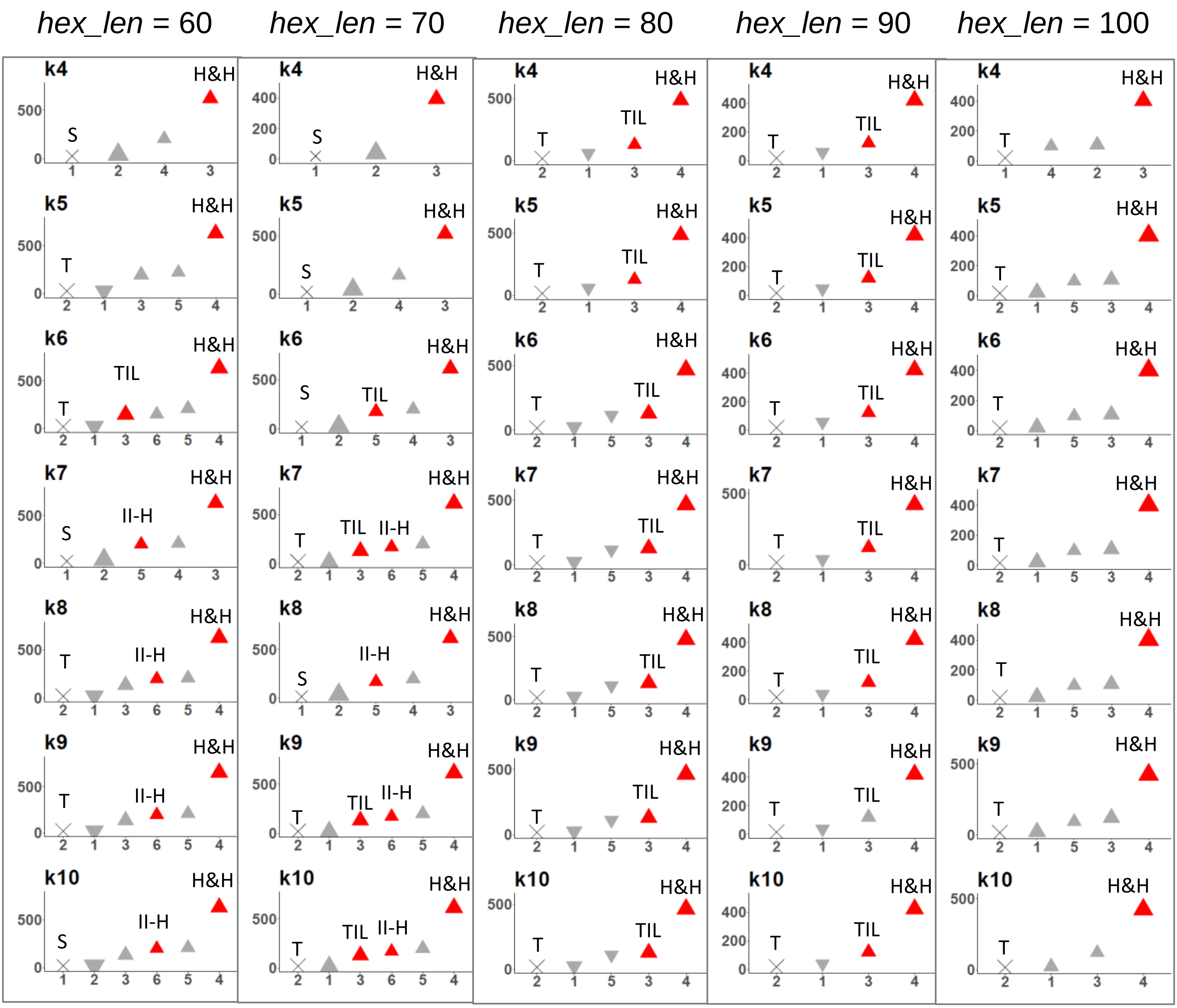


**Supplementary Figure 5.** Robustness analysis of TIPC using different sub-region sizes (hex_len) and number of clusters (k), based on CD3$^{+}$ T-cell CRC data. Prognostic significance based on univariate Cox PH regression model was used as the performance indicator. Vertical axis indicates the cluster-mean CD3$^{+}$ density; horizontal axis denotes the cluster identity. Clusters were ordered from left to right with from lowest to highest immune densities. The cluster with the lowest density was used as the reference in survival analysis and clusters exhibited significantly ($p < 0.05$) different survival outcome were color coded in red; $\Delta$ indicates better and $\nabla$ indicates worse survival than the reference; size of triangles represents the relative cluster size. Characteristic spatial subtypes : S = stroma-rich, T = tumor-rich, TIL = TIL-rich, II-H = I:T high-I:S high, H&H = hot-and-homogeneous.

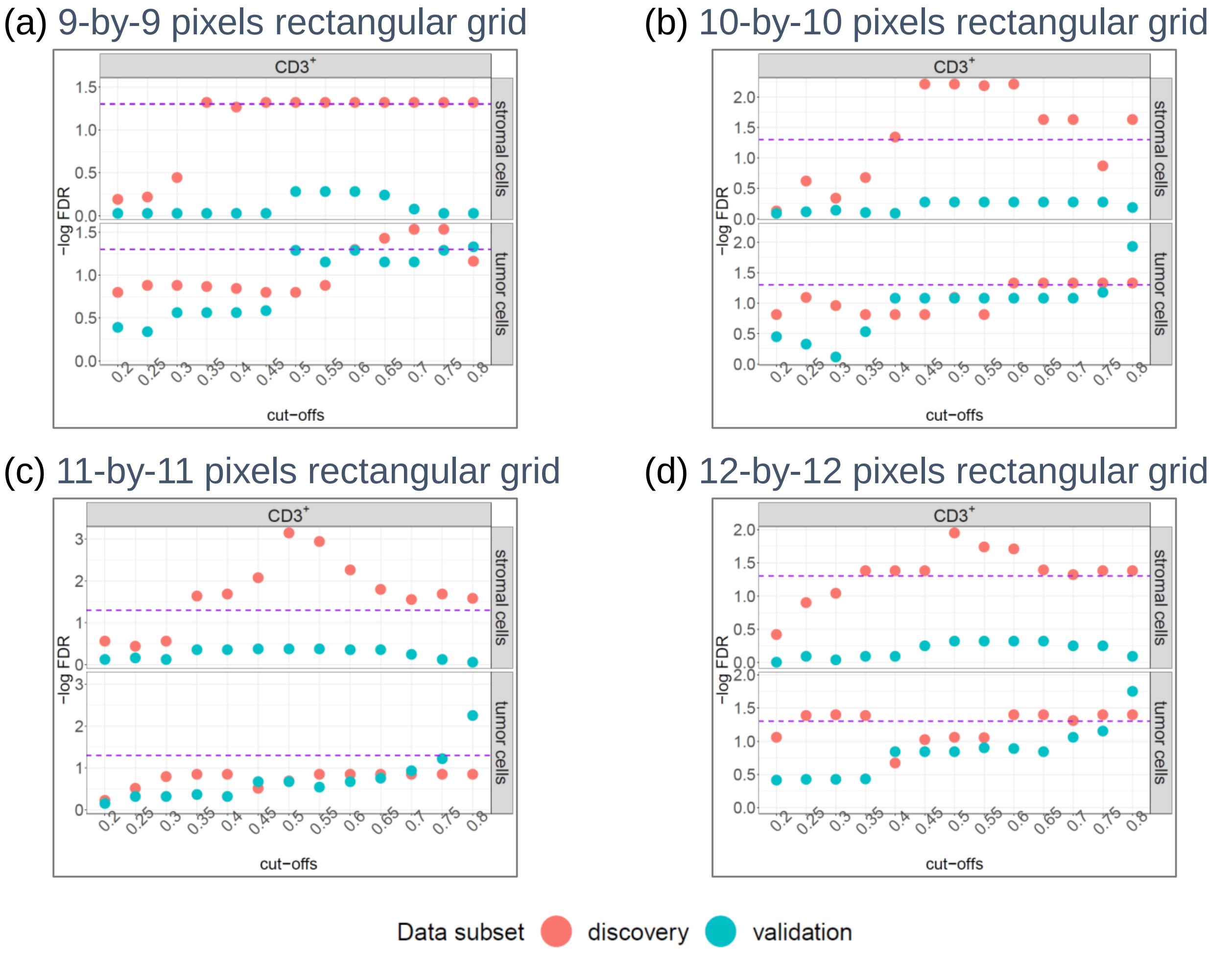


**Supplementary Figure 6.** Morisita-Horn (MH) analysis results using (**a**) 9-by-9, (**b**) 10-by-10, (**c**) 11-by-11, and (**d**) 12-by-12 pixels rectangular grids, based on CD3$^{+}$ T-cell data. FDR was obtained from univariate Cox PH regression analysis after adjusting for the 13 cut-offs as indicated on the horizontal axis. Upper panel: $MH_{stroma:CD3+T\ cell}$; lower panel: $MH_{tumor:CD3+T\ cell}$. Data points above the purple dotted lines (i.e. FDR < 0.05) are considered significant.

(a) 10-by-10 pixels rectangular grid

(b) 12-by-12 pixels rectangular grid

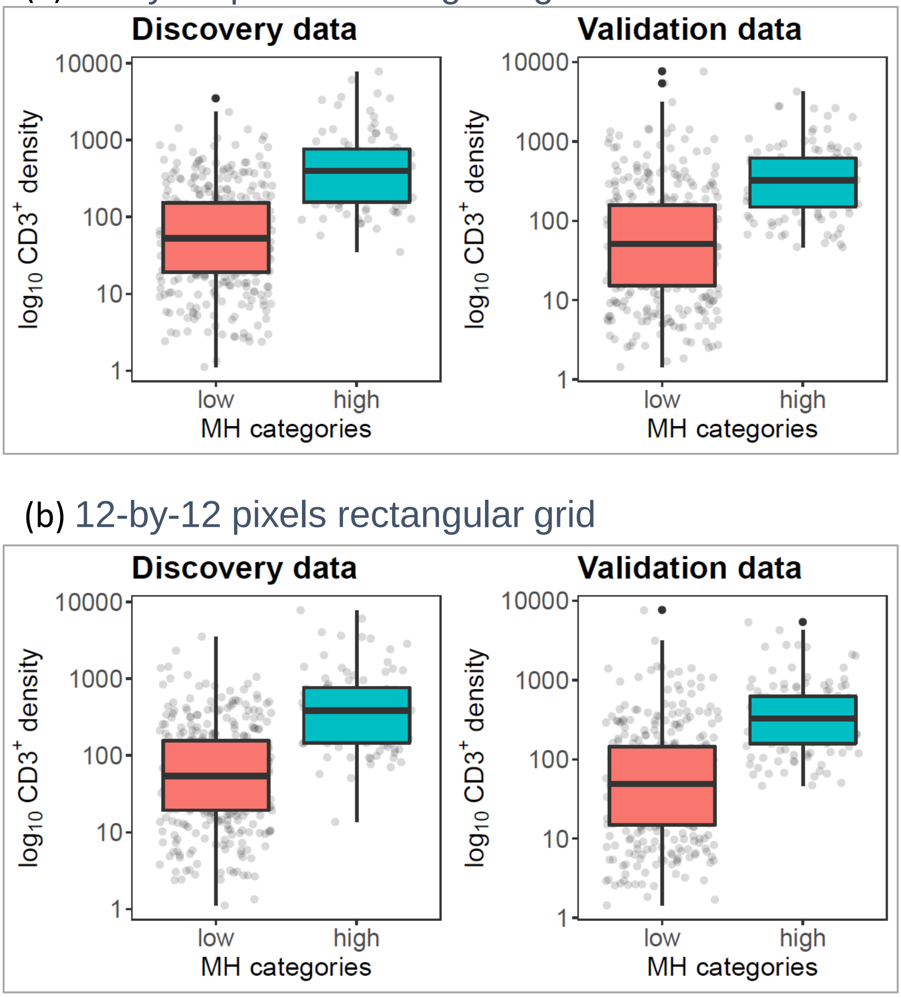


**Supplementary Figure 7.** Comparison of overall $CD3^+$ T-cell density distribution on Morisita-Horn (MH) dichotomized high and low groups (80th percentile was used as the cut-off); These MH analyses were performed using $CD3^+$ T cells and tumor cells, on (**a**) 10-by-10 and (**b**) 12-by-12 pixels rectangular grids. Box plot is defined by the 25th percentile (lower) and 75th percentile (upper) while extending lines mark the minimum (lower) and maximum densities (upper), black dots represent outliers higher (or lower) than the highest (or lowest) value within 1.5× the interquartile range (IQR); jittered gray dots represent cases.

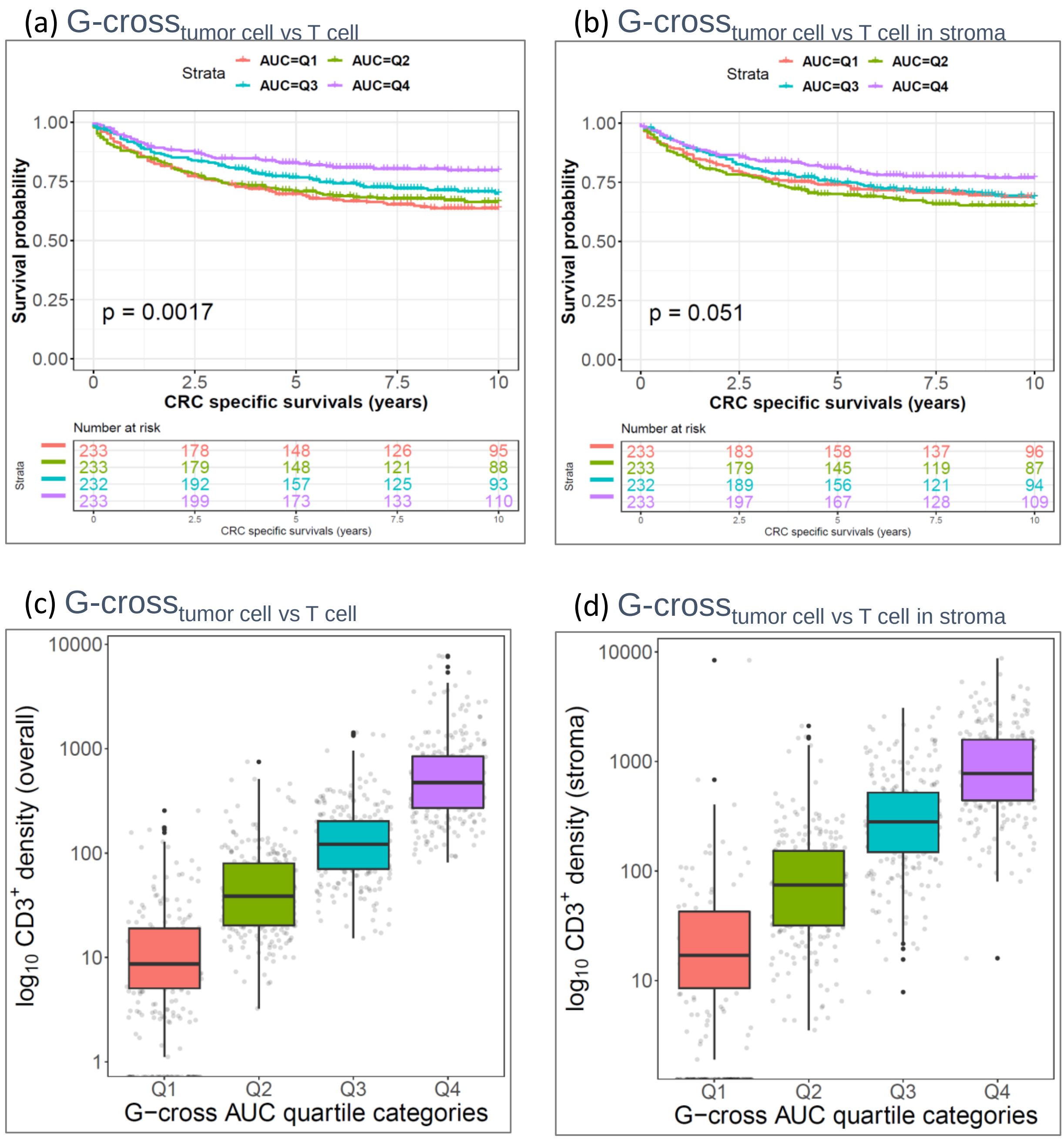


**Supplementary Figure 8.** G-cross function estimated (Kaplan-meier estimator) (**a**)(**c**) G-cross$_{\text{tumor cell vs T cell}}$ or (**b**)(**d**) G-cross$_{\text{tumor cell vs T cell in stroma}}$. (**a-b**) Kaplan-Meier curves and log-rank test p-values. Box-plots show (**c**) overall and (**d**) stromal CD3$^{+}$ T-cell density distribution across the tumor subtypes defined using G-cross AUC at 20μm. (**c-d**) Box plot is defined by the 25th percentile (lower) and 75th percentile (upper) while extending lines mark the minimum (lower) and maximum densities (upper), black dots represent outliers higher (or lower) than the highest (or lowest) value within 1.5× the interquartile range (IQR); jittered gray dots represent cases.

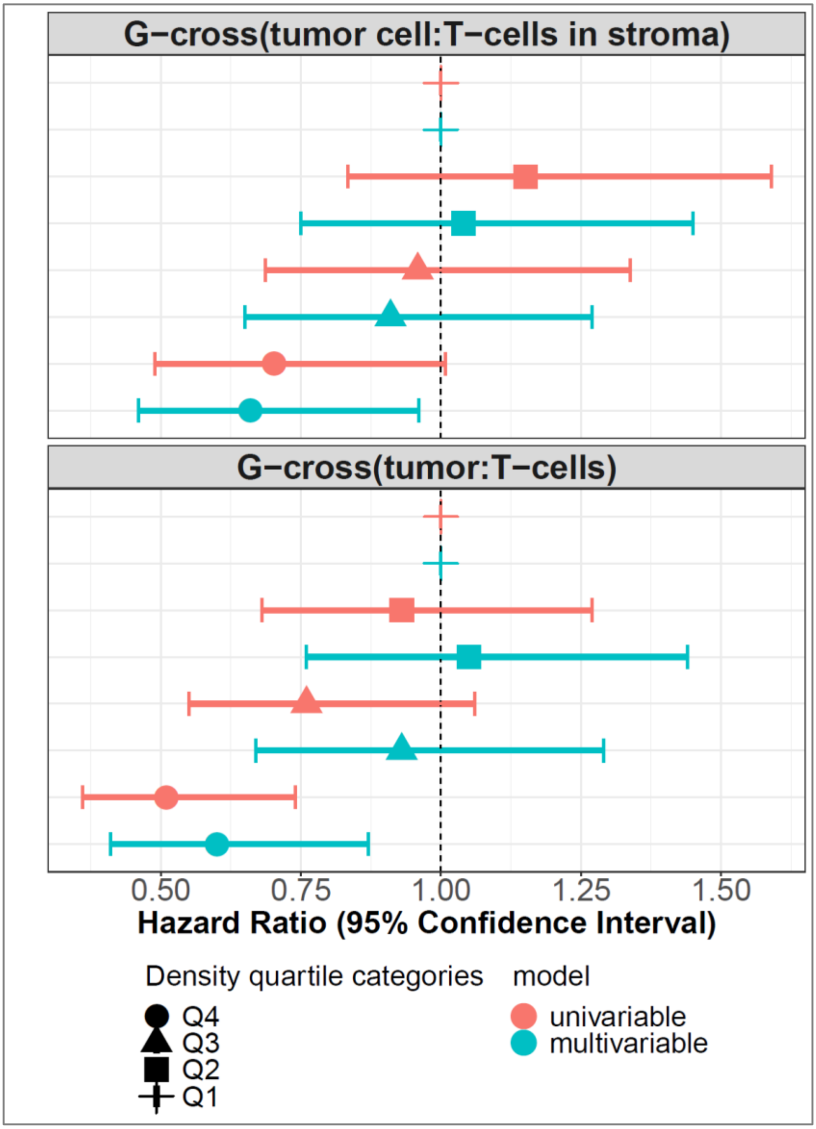


**Supplementary Figure 9.** Forest plots of HRs obtained from cox proportional hazards analyses using tumor subtypes determined using G-cross AUC; top panel: G-cross$_{\text{tumor cell vs T cell}}$, bottom panel: G-cross$_{\text{tumor cell vs T cell in stroma}}$; Q1: tumors with lowest quartile of G-cross AUC, Q4: tumors with highest quartile of G-cross AUC. The thin horizontal lines i.e. whiskers indicate the magnitude of the confidence interval: lower 95% (left) and upper 95% (right).

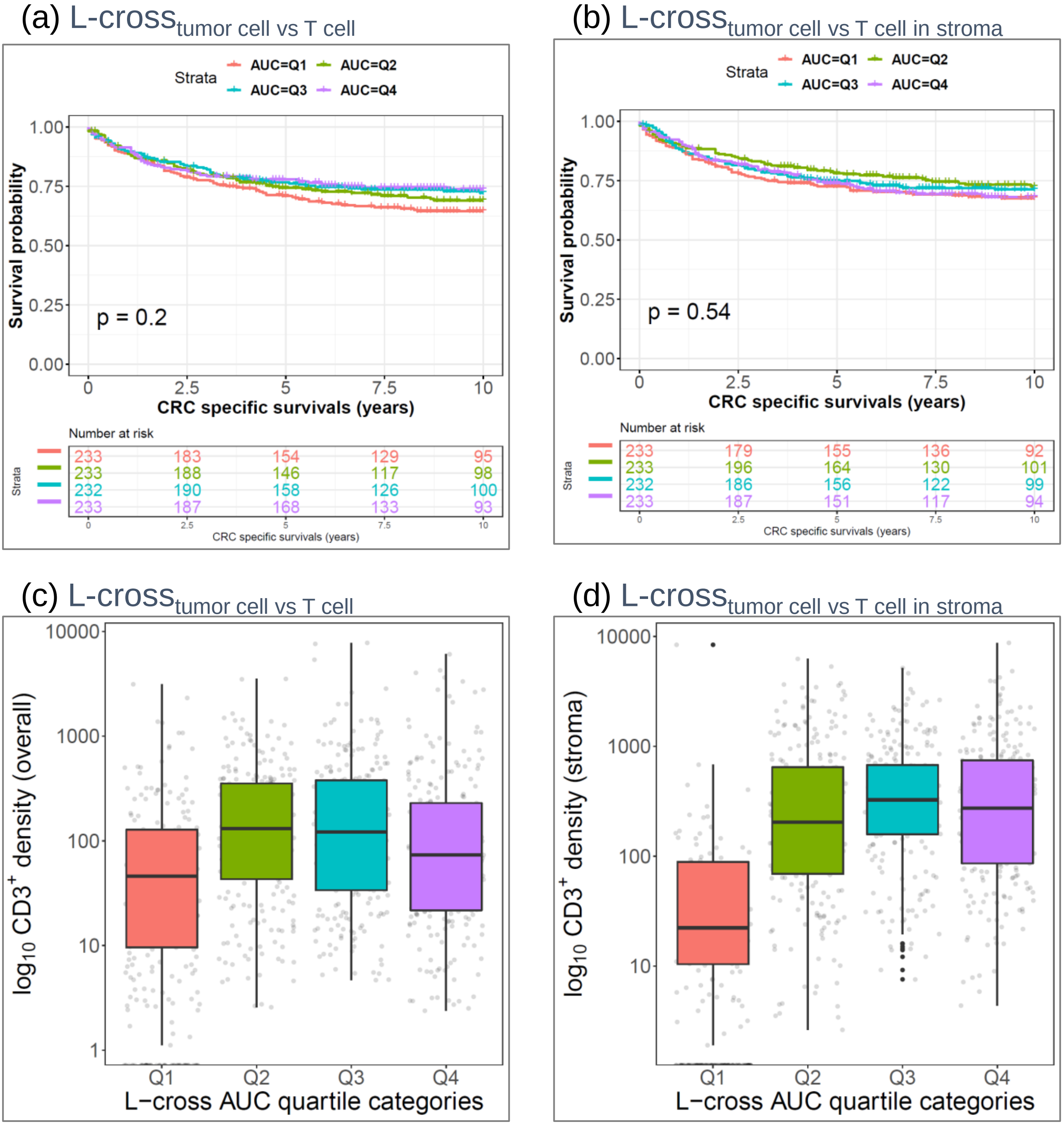


**Supplementary Figure 10.** L-cross function-based analyses (border estimator) (**a**)(**c**) L-cross$_{\text{tumor cell vs T cell}}$ or (**b**)(**d**) L-cross$_{\text{tumor cell vs T cell in stroma}}$. (**a**-**b**) Kaplan-Meier curves and log-rank test p-values. Box-plots show (**c**) overall and (**d**) stromal $CD3^+$ T-cell density distribution across the tumor subtypes defined using L-cross AUC. (**c**-**d**) Box plot is defined by the 25th percentile (lower) and 75th percentile (upper) while extending lines mark the minimum (lower) and maximum densities (upper), black dots represent outliers higher (or lower) than the highest (or lowest) value within 1.5× the interquartile range (IQR); jittered gray dots represent cases.

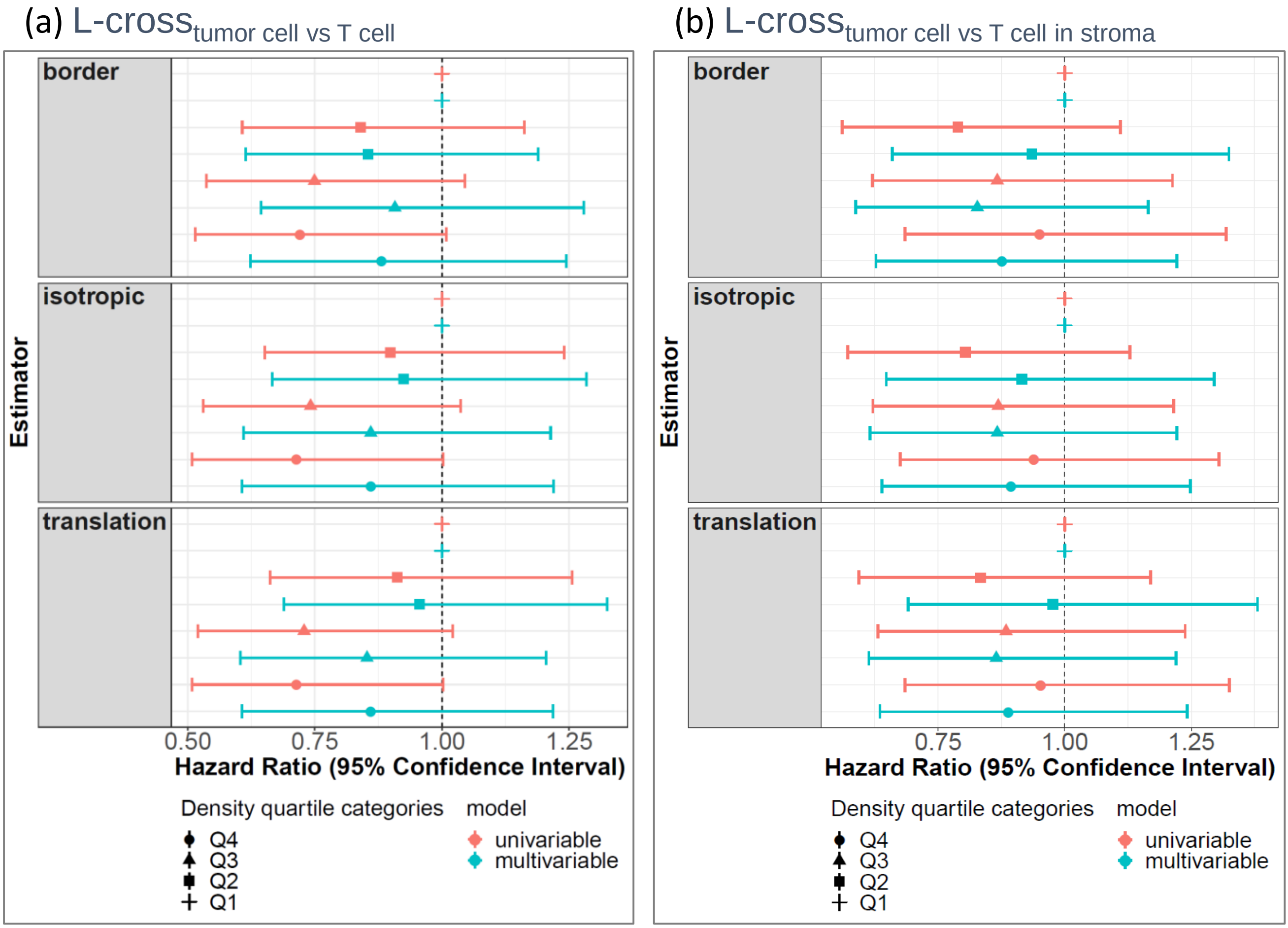


**Supplementary Figure 11.** Forest plots of HRs obtained from cox proportional hazards analyses using tumor subtypes determined using L-cross AUC (**a**) L-cross$_{\text{tumor cell vs T cell}}$ or (**b**) L-cross$_{\text{tumor cell vs T cell in stroma}}$, using cox proportional hazards analyses; Q1: tumors with lowest quartile of L-cross AUC, Q4: tumors with highest quartile of L-cross AUC. The thin horizontal lines i.e. whiskers indicate the magnitude of the confidence interval: lower 95% (left) and upper 95% (right).

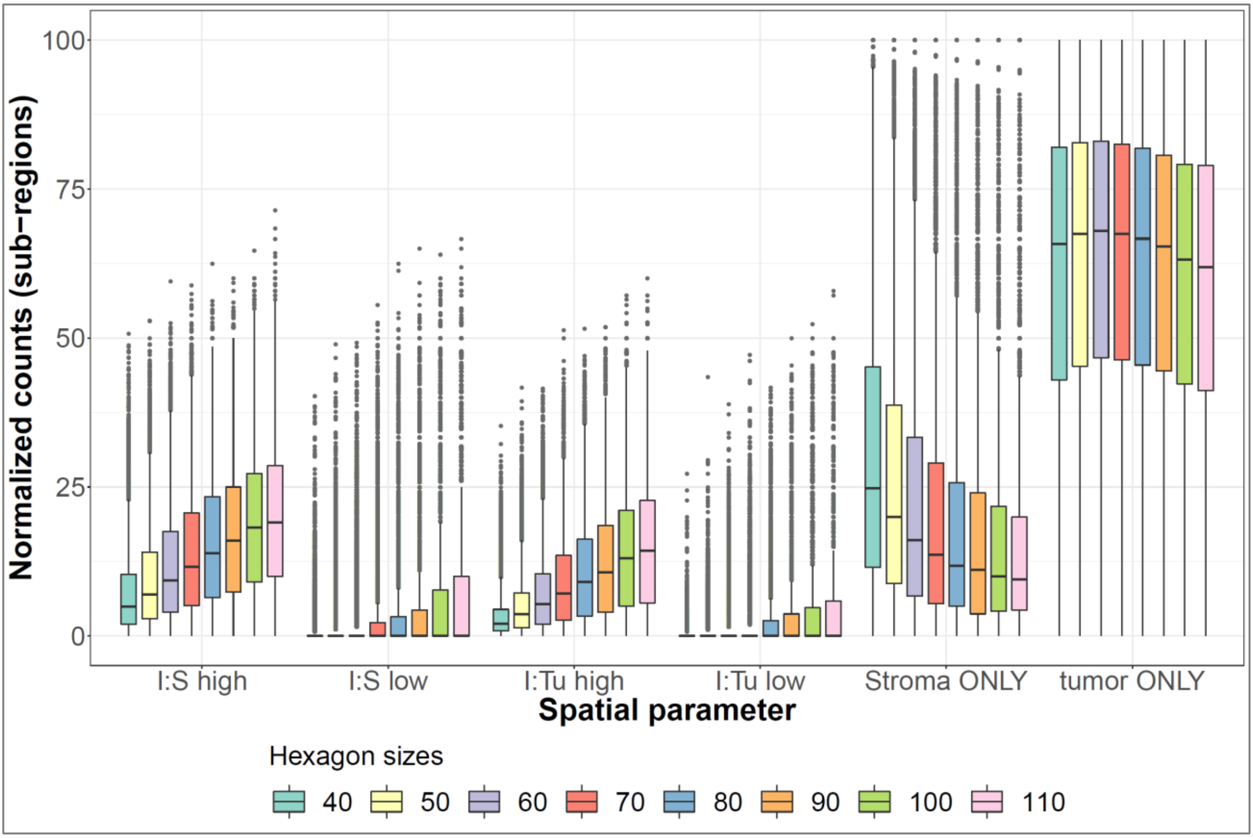


**Supplementary Figure 12.** Trend plot shows the distribution of %composition of the six TIPC spatial metrics at different sub-region sizes (i.e. hex_len) based on neutrophil CRC data. Box plot is defined by the 25th percentile (lower) and 75th percentile (upper) while extending lines mark the minimum (lower) and maximum densities (upper); gray dots represent cases.

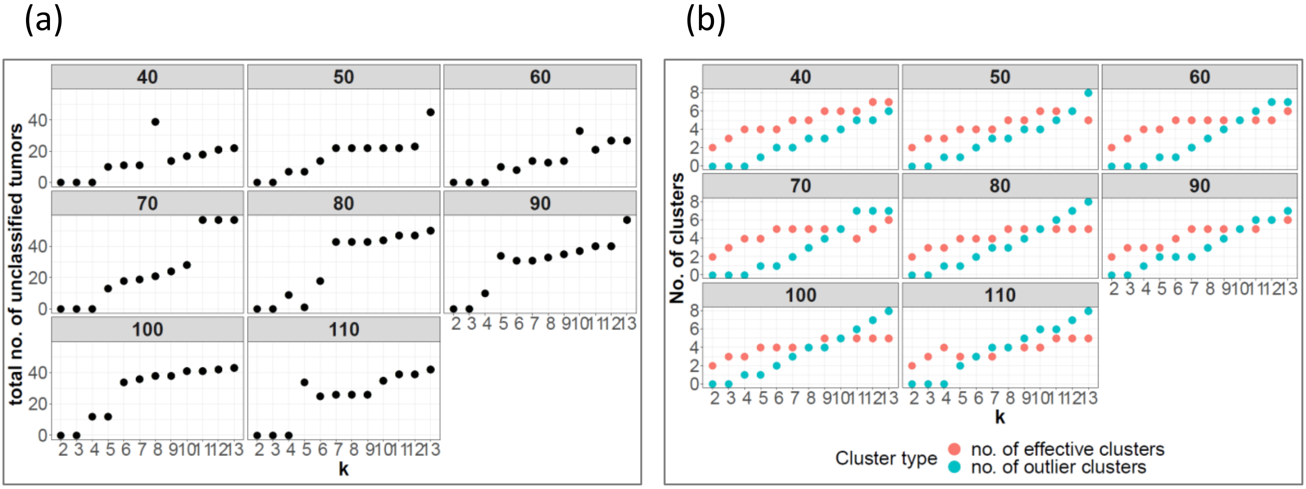


**Supplementary Figure 13.** Trend plots show (**a**) the total number of unclassified tumors i.e. those were detected in outlier clusters, (**b**) the number of outlier and effective (i.e. major clusters) clusters, obtained at different sub-region sizes (*hex_len* = 40 to 110, at interval of 10 pixels) and number of clusters (*k*). TIPC analysis was conducted using neutrophil CRC data. Outlier clusters were defined as clusters comprised of fewer than 30 tumors.

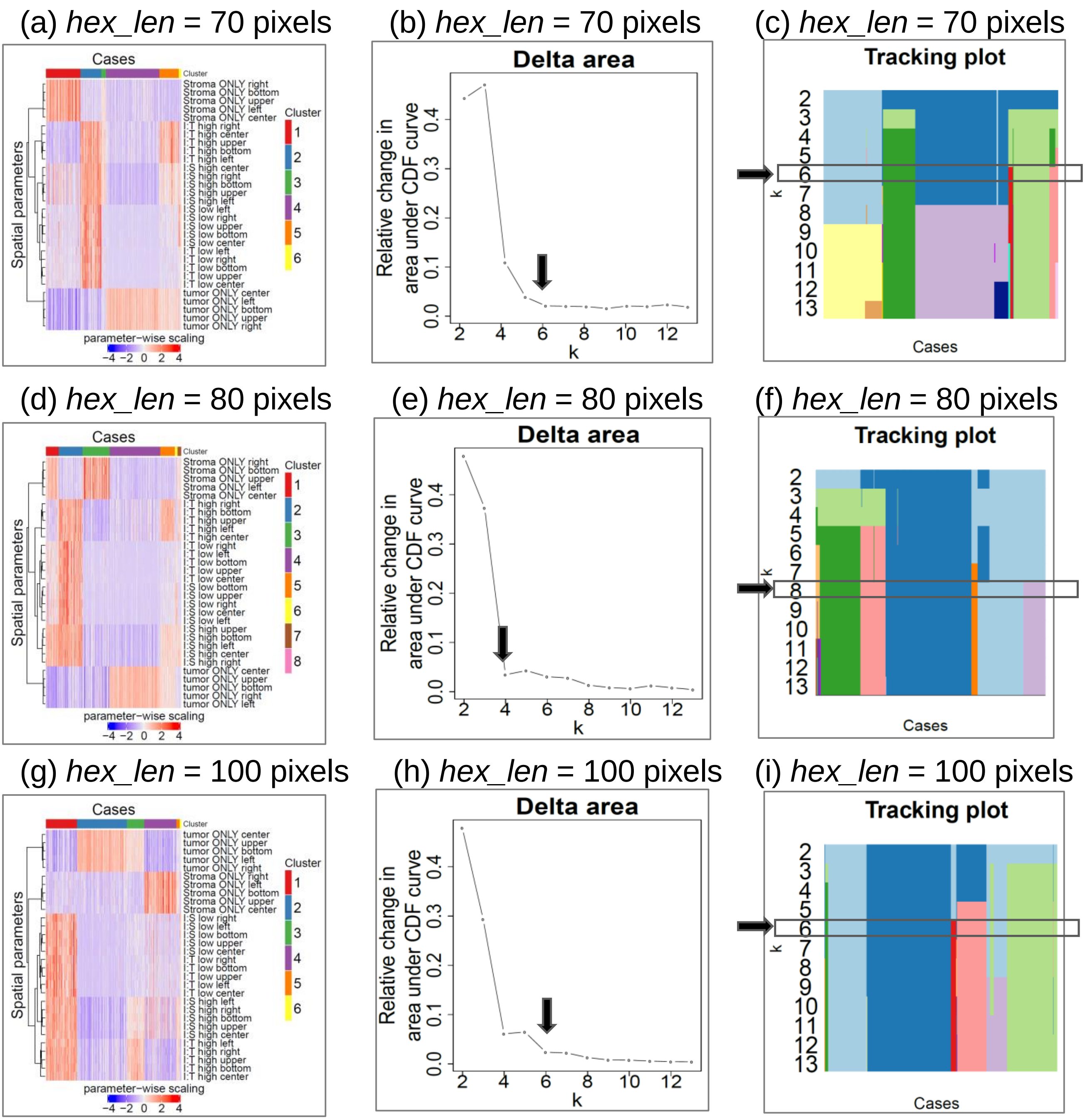


**Supplementary Figure 14.** Comparison of 3 different sub-region sizes, (**a**-**c**) 70, (**d**-**f**) 80, and (**g**-**i**) 100 pixels, used in TIPC application to neutrophil CRC data. The resulting hierarchical clustering results are shown in (**a**,**d**,**g**), and the corresponding auxiliary plots are displayed in (**b**,**e**,**h**) consensus cumulative distribution function (CDF) delta plots, and (**c**,**f**,**i**) tracking plots based on which the optimal number of clusters (*k*) was chosen.

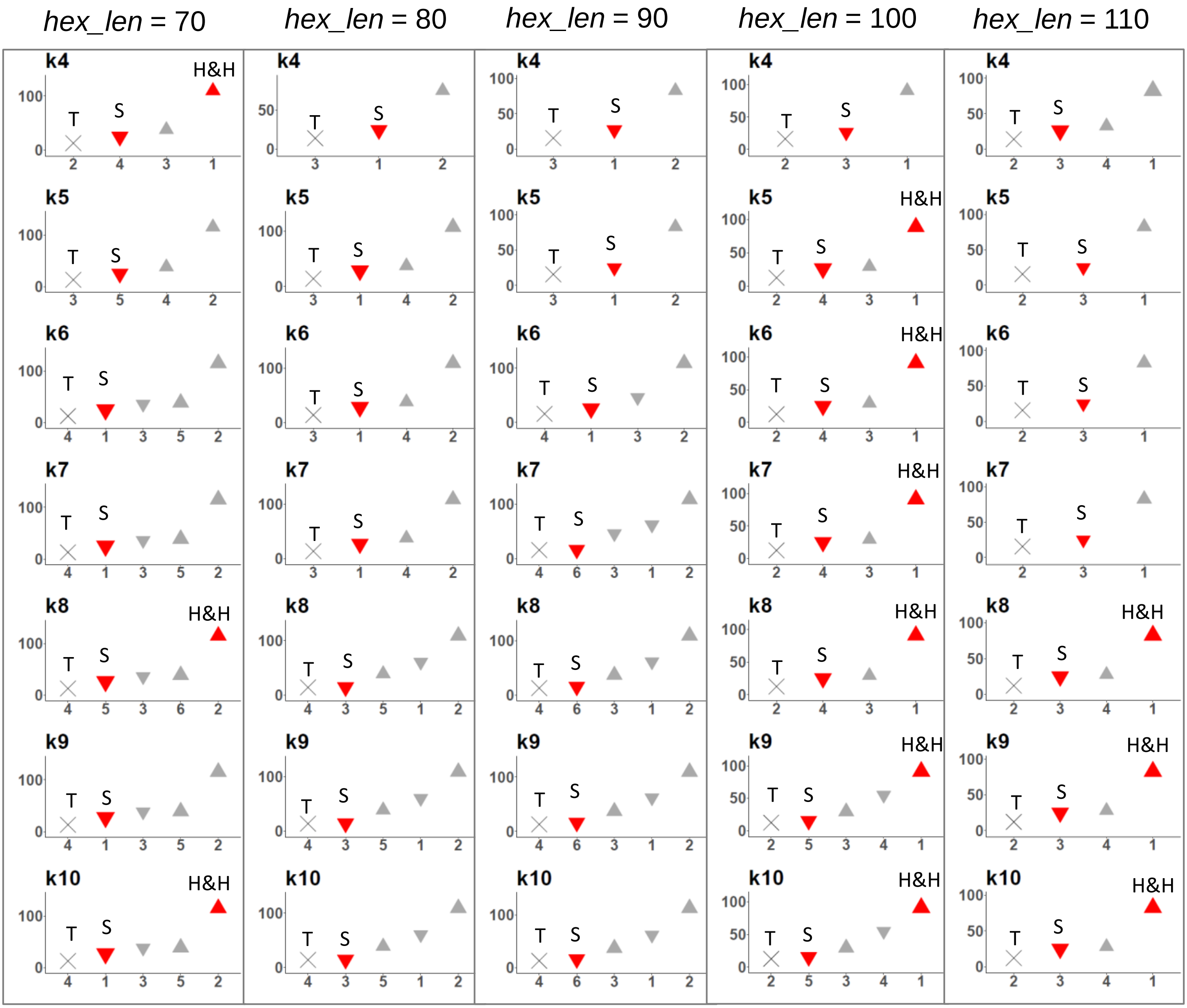


**Supplementary Figure 15.** Robustness analysis of TIPC using different sub-region sizes (*hex_len*) and number of clusters (*k*), using neutrophil CRC data. Prognostic significance based on univariate Cox PH regression model was used as the performance indicator. Vertical axis indicates the cluster-mean neutrophil density; horizontal axis denotes the cluster identity. Clusters were ordered from left to right with from lowest to highest immune densities. The cluster with the lowest density was used as the reference in survival analysis and clusters exhibited significantly ($p < 0.05$) different survival outcome were color coded in red; ∆ indicates better and ∇ indicates worse survival than the reference; size of triangles represents the relative cluster size. Characteristic spatial subtypes: S = stroma-rich, T = tumor-rich, H&H = hot-and-homogeneous.

Supplementary Table 1. Definition of the use of 'spatial' terminology

| Glossary of terms | Definition |
|---|---|
| Immune cell spatial organization or distribution | The relative spatial locations (i.e. XY coordinates in a two dimensional Cartesian coordinate system) of a collection of immune cells of interest in the TME. |
| Tumor-immune spatial relationship | The spatial distribution between immune cells and tumor (or stromal cells), at a smaller spatial scale, e.g. a dense immune cell cluster is found in near proximity to tumor cells (or stromal cells). |
| Tumor-immune spatial pattern (abbreviated as spatial pattern) | Unique combinations of tumor-immune spatial relationships observed at a larger spatial scale such as tumor whole-slide or microarray section |
| Spatially informed tumor subtypes (abbreviated as spatial subtypes) | Tumor subsets with similar tumor-immune spatial patterns. |
| Spatial composition of a tumor-immune microenvironment | The constitution of a tumor microenvironment in terms of (1) tumor-immune spatial relationship, (2) immune cell abundance, and (3) tumor morphology. |
| TIPC spatial parameters/ measures | The six-element vector, which encapsulates the tumor-immune spatial relationship in a TME, represented as six normalized counts of sub-region categorization |
| Spatial Point Patterns | A ‘spatial point pattern’ is a dataset giving the observed spatial locations of things or events. |

Supplementary Table 2. Definition of the six spatial parameters of TIPC algorithm

| TIPC sub-region classification | Definition |
|---|---|
| tumor-only | Sub-regions containing only tumor cells |
| immune-to-tumor low (I:T low) | Sub-regions containing both immune and tumor cells, and the local immune to tumor ratio is smaller than the global ratio (i.e. total number of immune cells over total number of tumor cells, in the tissue section) |
| immune-to-tumor high (I:T high) | Sub-regions containing both immune and tumor cells, and the local immune to tumor ratio is larger than the global ratio (i.e. total number of immune cells over total number of tumor cells, in the tissue section) |
| stroma-only | Sub-regions containing only stromal cells |
| immune-to-stroma low (I:S low) | Sub-regions containing both immune and stromal cells, and the local immune to stroma ratio is smaller than the global ratio (i.e. total number of immune cells over total number of stromal cells, in the tissue section) |
| immune-to-stroma high (I:S high) | Sub-regions containing both immune and stromal cells, and the local immune to stroma ratio is larger than the global ratio (i.e. total number of immune cells over total number of stromal cells, in the tissue section) |

Supplementary Table 3. Descriptive name definition for unique spatial patterns observed in this study, using TIPC

| Unique spatial subtypes (abbreviation) | Descriptions |
|---|---|
| hot-and-homogenous (H&H) | Tumor enriched in high proportions of sub-regions categorized under I:T high and low, and I:S high and low |
| I:T high-I:S high (II-H) | Tumor enriched in high proportions of sub-regions categorized under I:T high and I:S high; might be accompanied by tumor-ONLY or stroma-ONLY |
| tumor-rich (T) | Tumor enriched in high proportions of sub-regions categorized under tumor-only |
| stroma-rich (S) | Tumor enriched in high proportions of sub-regions categorized under stroma-only |
| TIL-rich | Tumor enriched in high proportions of sub-regions categorized under tumor-only as well as I:T high |
| SIL-rich | Tumor enriched in high proportions of sub-regions categorized under stroma-only and I:S high |
